# Does transection severity determine the way of spinal cord repair in the spiny mouse?

**DOI:** 10.64898/2026.07.17.739224

**Authors:** Aleksandr Veshchitskii, Aleksandr Mikhalkin, Polina Shkorbatova, Oleg Gorskii, Anton Beljajev, Olja Mijanović, Maria E. Velizhanina, Eduard Sharapenkov, Aleksandr A. Rubel, Natalia Merkulyeva

## Abstract

The mechanisms of the spinal cord regeneration after complete spinal cord transection were investigated in spiny mice. In some animals, the appearance of quadrupedal overground stepping together with rewiring of the direct propriospinal projections between the cervical and lumbar enlargements was revealed. In others, no stepping recovery was detected, whereas numerous cells labeled by the neuronal proteins NeuN and βIII-tubulin were observed within the injury region. We suggest that depending on trauma severity, different repair mechanisms are elicited: only connectome restoration or both connectome restoration and the activation of neurogenesis. To confirm the high neurogenic potential of spiny mice, a primary culture of bone marrow was established. Unlike in other mammals, bone marrow pluripotent cells in the culture differentiated into neuronal cells without any chemical stimulation. These findings provide strong evidence for the high differentiation potential of spiny mouse stem cells toward neural lineages.

**GRAPHICAL ABSTRACT:** 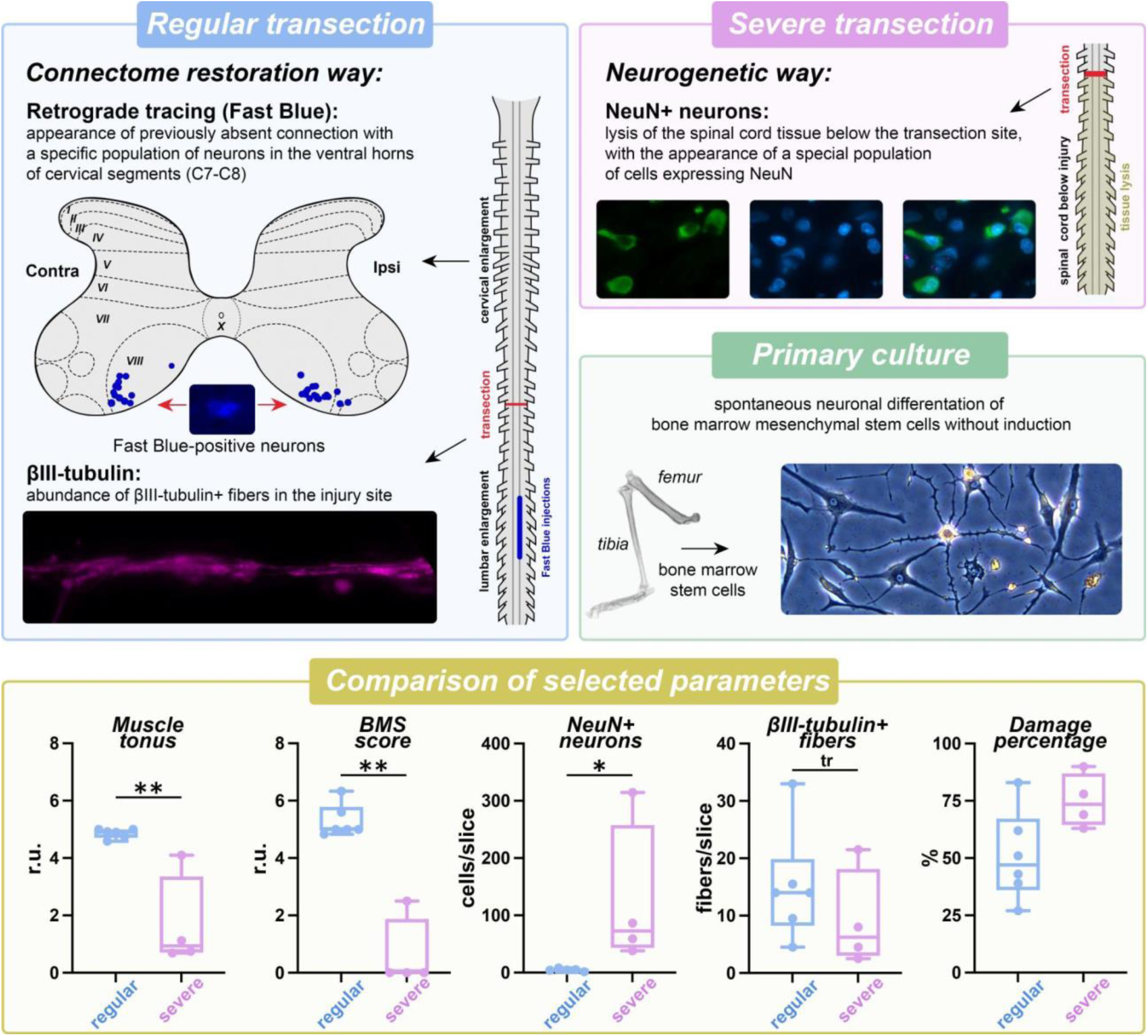

**HIGHLIGHTS:**

- Two regenerative mechanisms are proposed in spiny mice, depending on the severity the spinal cord transection
- Regular transection evoked the emergence of the direct propriospinal projections
- Severe transection evoked the neurogenesis within the primary injured region
- Primary culture of bone marrow cells from spiny mice exhibits neurogenic differentiation without chemical induction

## INTRODUCTION

Traumatic central nervous system (CNS) damage is a devastating condition associated with high morbidity and mortality. Traumatic brain injury affects 50-60 million people each year (Maas et al., 2022). The incidence of traumatic spinal cord injury ranges from 3.3 to 195.4 cases *per* million (Jazayeri et al., 2023). At the same time, traumatic CNS damage is among the least repairable injuries. One of the main reasons for this is the formation of a glial or fibrotic scar within the injured tissue (Tran et al., 2022; Clifford et al., 2023), preventing adequate axonal sprouting and cell migration.

Alongside multiple rehabilitation techniques based on the external activation of the injured spinal networks (see Ko, 2023), new animal models with unique regenerative abilities may help in the development of new strategies to trigger inherent repair mechanisms. One such model is the spiny mouse from the family *Muridae*, which exhibits a well-developed ability for scar-free regeneration of the skin (Stewart et al., 2018; Yoon et al., 2020) and muscles (Maden et al., 2018; Peng et al., 2021). Recently, evidence supporting scar-free regeneration of the spiny mouse CNS has also been provided (Streeter et al., 2020; Nogueira-Rodrigues et al., 2022).

At the same time, the mechanisms underlying such scar-free CNS regeneration in spiny mice remain to be discovered. Meanwhile, several key findings have already been reported: reduced spinal inflammation and fibrosis (Streeter et al., 2020), enhancement of axon regeneration (Nogueira-Rodrigues et al., 2022), and inter-neuronal connectome reorganization (Kidd et al., 2024). Notably, alongside evidence for connection restoration, an absence of recovery of the nervous tissue cytoarchitectonic organization has been documented (Nogueira-Rodrigues et al., 2022; Kidd et al., 2024). These findings suggest that CNS regeneration in spiny mice may be based on separate mechanisms with unequal effectiveness: those of axonal regrowth/remodelling and those of neurogenesis/cytoarchitectonic repair. Indeed, in mammals, axonal sprouting is one of the mechanisms of CNS regeneration (Massey et al., 2006; Rossignol et al., 2009; Bachmann et al., 2014); and *de novo* neuronal proliferation has also been documented after brain and spinal cord injury (Jin et al., 2001; Ke et al., 2006; Vessal et al., 2007; McDonough et al., 2013). Therefore, the aim of the present study was investigation of the mechanisms of the functional and structural recovery in spiny mice after complete spinal cord transection.

Functional recovery was assessed using: (1) the Basso, Beattie, Bresnahan (BBB) locomotor rating scale (Basso et al., 1995) and Basso mouse scale (BMS) (Basso et al., 2006), and (2) analysis of overground locomotion. Histological recovery in the injury site was assessed by analysing: (1) the distribution patterns of the following cytoskeleton components: βIII-tubulin, which labels neurite outgrowth profiles (Wang et al., 2018); (2) direct descending projections from the supraspinal regions and the upper spinal cord across the injury site to the lumbar segments using a retrograde tracer Fast Blue (FB); and (3) the general neuronal population using the neuronal protein NeuN (Mullen et al., 1992). We also assessed the levels of neuroinflammation and astrogliosis using selective macrophage/microglial (ionized Ca^2+^-binding adapted molecule 1, Iba1) and astroglial (glial fibrillary acidic protein, GFAP) markers (see Freyermuth-Trujillo et al., 2022; Bhatt et al., 2024).

## RESULTS

## 1. Injury assessment

The first task was to assess the completeness of the spinal cord transection (**Figure 2E, upper right panels)**. As described in the Methods, two types of transection were performed: using the micro-scissors and the microscalpel (complete transection 1, *TRC1* group) and using the microscalpel and the hook (complete transection 2, *TRC2* group) (**Figure S1**). In the *TRC1* group in 1/7 animals, the transection was identified as incomplete: bilateral ventral-most regions composing 11% of the transverse spinal cord slice remained intact. Hereafter, this animal is referred to as the *TRI* animal (incomplete transection). In the *TRC2* group, no incomplete transection was identified.

We quantitatively assessed the damaged areas within the spinal cord region extending 2 mm caudally and 2 mm rostrally to the primary injury site and calculated the ratio of damaged area to the total region of interest. The assessed zone was divided into two subzones: 0-1 mm and 1-2 mm from the primary injury (**Figure 2E, upper right panels**). The *TRI* animal had a comparatively low damage rate: in the 1-2 mm and 0-1 mm caudal zones, damage was 0% and 52%, respectively; in the 1-2 mm and 0-1 mm rostral zones, damage was 0% and 49%, respectively; the average total damage was 28%. In the *TRC1* group, the damage pattern was largely symmetric: caudal damage was 68 ± 25% in the 0-1 mm zone and 28 ± 26% in the 1-2 mm zone; rostral damage was 67% ± 21% in the 0-1 mm zone and 27 ± 29% in the 1-2 mm zone (**Figure 2E, lower right panels**). The least and the most total damage were observed in animals #*142* and *#137* (27% and 83%, respectively). In 3/4 animals from the *TRC2* group, the entire assessed caudal area was completely damaged, with no signs of remaining intact tissue (**Figure S1**). In 1/4 animals (#*146*), only the 0-1 mm caudal zone was almost fully damaged (94%), while the more distant 1-2 mm caudal zone contained largely intact tissue (39% damaged). Moreover, this animal had total damage comparable to that of the *TRC1* group (63% compared to 69-90% in other *TRC2* animals). Statistical comparisons revealed that the *TRC1* group had a significantly smaller caudal damaged area compared to the *TRC2* group (U = 2.0, p = 0.0333; M-W) but a similar rostral damaged area (U = 8.0, p = 0.4381; M-W). The comparison of total damage did not show a significant difference (*TRC1 vs TRC2*: U = 3, p = 0.0667).

In most animals, well-defined cystic cavities (empty or containing an amorphous component) were observed within the primary and secondary injury regions (**Figure 2E, upper right panels, and S1**). Similar cystic cavities were previously observed in the injured spinal cord of spiny mice (Nogueira-Rodrigues et al., 2022) and rats (Sroga et al., 2003; Telegin et al., 2023). Interestingly, a delayed activation of inflammatory cells was proposed to be a reason for the cavitation in rats (Sroga et al., 2003). However, a muted inflammatory response was observed in the skin of spiny mice (Ghebryal et al., 2023).

## 2. Sensorimotor functions

### 2.1. Muscle tonus

Before injury, a maximal score (“5”) for hindlimbs muscle tonus was documented in all animals. All injured animals exhibited decreased muscle tonus at the 1st week post-injury (WPI). In 2/6 animals from the *TRC1* group, the maximal muscle tonus score was achieved by the 6th WPI; in 2/6 animals, by the 7th WPI, and in the remaining 2/6 animals, no maximal score was reached, even by the 9th WPI (**Figure 1A**). In the *TRC2* group, only 1/4 animals achieved the maximal level of the muscle tonus *(#146)*. In the *TRI* animal, the maximal muscle tonus score was achieved by the 3th-5th WPI (**Figure 1A**). In the *Sham* group, the maximal muscle tonus score was observed by the 2nd-3rd WPI (**Figure 1A**). At the 6th WPI, significant differences between the *TRC1* and *TRC2* groups were revealed (U = 0, p = 0.0095; M-W) (**Figure 1A**).

**Figure 1.**
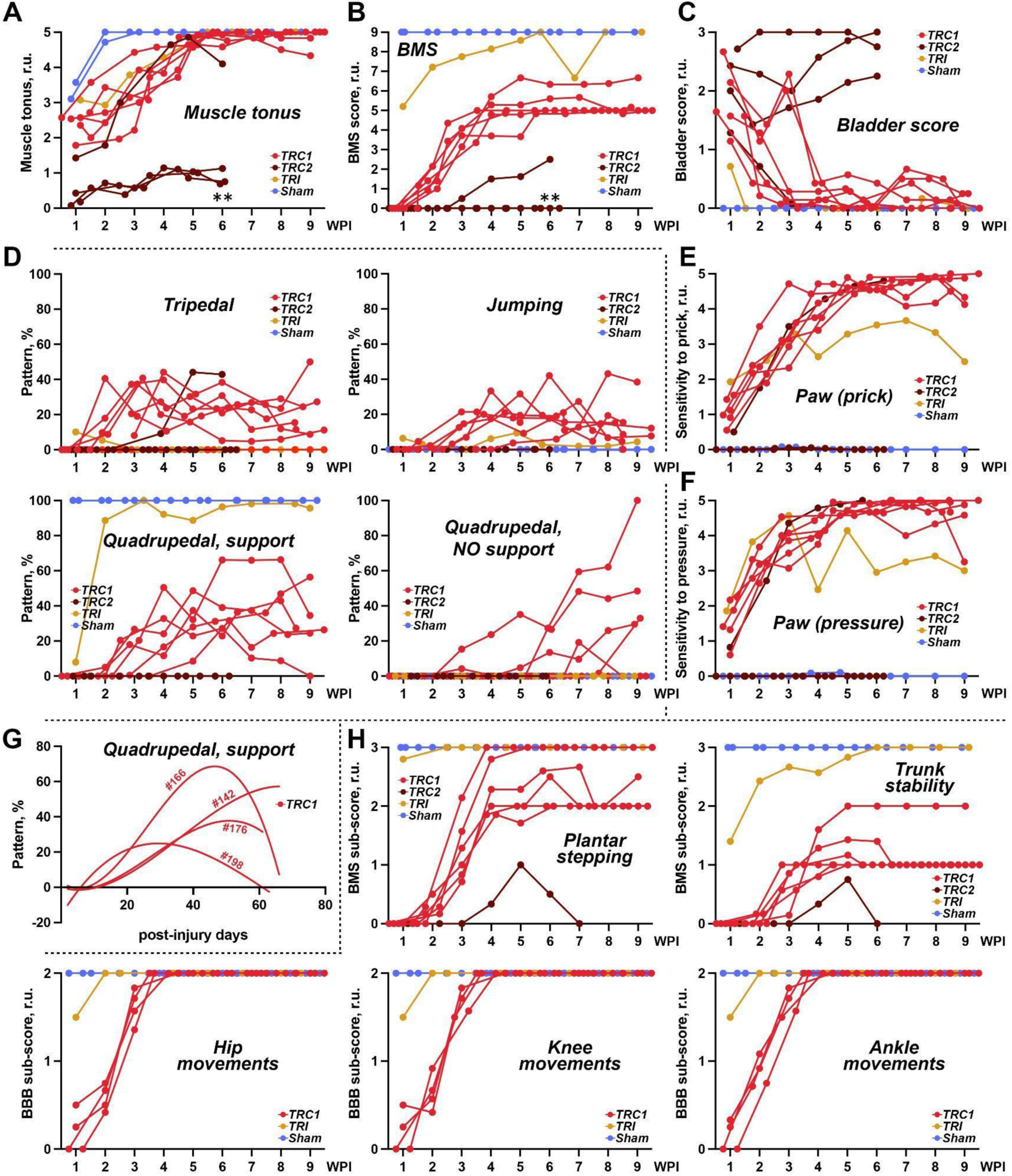
A post-injury dynamic of the physiological status after the complete (*TRC1* and *TRC2*, red and brown respectively) and incomplete (*TRI*, orange) transection, and in sham-operated animals (*Sham*, blue). (A) Muscle tonus (Mann-Whitney U test, ** – p < 0.001). (B) BMS score (Mann-Whitney U test, ** – p < 0.001). (C) Bladder tonus and the dominated size of the urine spots. (D) Post-injury dynamic of four stepping patterns: tripedal, jumping, quadrupedal stepping with support, and quadrupedal stepping without support. (E) Hypersensitivity to the paw pinprick. (F) Hypersensitivity to the paw pressure. (G) Examples of the post-injury dynamic of the percent occurrence of the quadrupedal stepping with weight support in four animals from the *TRC1* group (#142, #166, #176, and #198). Solid lines are envelope curves. (H) A post-injury dynamic of some parameters of the BMS (plantar stepping and trunk stability) and BBB sub-scores (hip, knee, and ankle movements). For all panels, WPI – weeks post-injury.

### 2.2. BMS score

Before the injury, the BMS score in all animals corresponded to the maximal level (“9”). In *TRC1* animals, the BMS score at the 1st WPI was at the zero level. Over time, an improvement in the BMS score was observed, however, the highest BMS score in the *TRC1* group did not exceed the “5-6” level (**Figure 1B**). In the *TRC2* group, only one animal (#*146*) achieved the BMS score higher than the zero level (1.5-2.5) (**Figure 1B**). The *TRI* animal reached the maximal BMS score (“9”) by the 5th WPI (**Figure 1B**). *Sham* animals maintained the maximal BMS score throughout the entire postoperative period (**Figure 1B**). At the 6th WPI, significant differences between the *TRC1* and *TRC2* groups were observed (U = 0, p = 0.0048; M-W) (**Figure 1B**).

### 2.3. Locomotor patterns

Overground locomotion was assessed beginning from the 1st WPI. In all *TRC1* animals, the single locomotor pattern during the 1st WPI was *bipedal* locomotion using the forelimbs (**Video S1**). This pattern decreased sharply over time, remaining mainly below 15% of all stepping episodes. During the 2nd WPI, all animals also exhibited *tripedal* locomotion (**Video S1**). The tripedal pattern increased sharply by the 2nd WPI and remained at a level of 15-30% throughout the post-injury period (**Figure 1D, upper left panel**). At the 3rd WPI, all animals also exhibited a *tripedal pattern with a shift* in which the hindlimb was placed later behind the stride. This pattern was quite stable (around 10%) throughout the entire post-injury period. At the 2nd WPI, two animals also performed a *jumping* pattern with weight loading on the forelimbs (**Video S1**); three other animals performed it at the 3rd WPI, and one animal performed it at the 4th WPI (**Figure 1D, upper right panel**). The jumping pattern increased sharply by the 4th WPI and remained at a level of 15% throughout the post-injury period (**Figure 1D, upper right panel**). *Quadrupedal stepping with support* (**Video S1**) was observed in one animal at the 2nd WPI, in four animals at the 3rd WPI, and in one animal at the 5th WPI (**Figure 1D, lower right panel**). This pattern increased sharply by the 4th WPI and remained at a level of 35% throughout the post-injury period (**Figure 1D, lower right panel**).

Starting from the 3rd-8th WPI, 5/6 animals from the *TRC1* group also performed *quadrupedal stepping without support* (**Video S1**). The emergence of this pattern is the main reason why, starting from the 5th-7th WPI, quadrupedal locomotion with support gradually decreased (**Figure 1G**). We named this phenomenon “*a secondary disruption of stepping*”. Only one animal from the *TRC1* group (#*142*) did not exhibit quadrupedal locomotion without support. To better illustrate the dynamics of quadrupedal stepping with support, cubic regression was performed (**Figure 1G**). For this purpose, daily data (not weekly averages) were used. Animals with the strongest decrease (#*166* and #*198*), intermediate decrease (#*176*), and no decrease (#*142*) were compared.

Notably, quadrupedal locomotion without support and quadrupedal locomotion with support were easily intermixed within the same walking episode. In many cases, jumping episodes were observed just before the transition between the patterns without and with support.

In the *TRC2* group, 3/4 animals performed only bipedal stepping throughout the entire postoperative period. Only one animal (*#146*) was able to perform tripedal locomotion with partial weight support, starting at the boundary between the 3rd and 4th WPI. The occurrence of this pattern increased from 9.2% at the 4th WPI to 42-44% at the 5th and 6th WPI (**Figure 1D, upper left panel**).

The *TRI* animal exhibited early onset of quadrupedal stepping with support: at the 1st-2nd WPI (**Figure 1D, lower left panel**). The percentage of this pattern reached 100% at the 3rd WPI. This animal did not exhibit quadrupedal stepping without support (**Figure 1D, lower right panel**). The jumping pattern was observed in this animal throughout the entire post-injury period (**Figure 1D, upper right panel**).

All *Sham* animals exhibited exclusively quadrupedal locomotion with support throughout the entire postoperative period (**Figure 1D, lower left panel**).

### 2.4. BMS and BBB sub-scores

Because over time a secondary disruption of stepping was detected in most *TRC1* animals, for some animals (*TRC1*: n = 6, *TRC2*: n = 1, *TRI*: n = 1, *Sham*: n = 2), we separately analysed sub-scores of the BMS and BBB tests related to leg movement (plantar stepping score and intensity of hindlimb movements in the hip, knee, and ankle joints) and to weight support (trunk stability) (**Figure 1H**). The plantar stepping score ranged from “0” (no plantar stepping) to “3” (consistent plantar stepping), the leg movement from “0” (no movement) to “2” (extensive movement), and the trunk stability from “0” (no stability) to “3” (full stability). Notably, because most animals in the *TRC2* group did not show BBB improvement, only data from the single animal (#*146*) were analysed.

#### Plantar stepping

In 2/6 *TRC1* animals, the plantar stepping score reached its maximal value (“3”) at the 4th WPI; in the others, it did not exceed the “2.5” value and maintained this level throughout the remaining post-injury period (**Figure 1H, upper left panel**). In the #*146* animal (*TRC2* group), a non-zero level of plantar stepping was observed only at the 4th-6th WPI (**Figure 1H, upper left panel**). *Sham* animals maintained the maximal plantar stepping score (“3”) throughout the entire postoperative period (**Figure 1H, upper left panel**). The *TRI* animal maintained the maximal plantar stepping score (“3”) throughout the entire postoperative period, except for a slight decrease at the 1st WPI (**Figure 1H, upper left panel**).

#### Leg movements

In the *TRC1* group, all leg movements scores (including hip, knee, and ankle movements) reached their maximal level at the 4th WPI (**Figure 1H, lower panels)**. Since leg movements were completely absent in the *TRC2* group (except for #146), it was not possible to assess BBB leg movements in these animals. In the *Sham* group, leg movements scores reached their maximal value during the 1st WPI (**Figure 1H, lower panels)**. In the *TRI* animal, leg movements scores reached their maximal value at the 2nd WPI (**Figure 1H, lower panels**).

#### Trunk stability

Just after the surgery, zero trunk stability was documented for the *TRC1* group (**Figure 1H, upper right panel**). By the 4th WPI, it reached its highest value but never achieved the maximal score (“3”) (**Figure 1H, upper right panel**). In the #*146* animal (*TRC2* group), a non-zero level of trunk stability was observed at the 4th-5th WPI; however, it declined by the 6th WPI (**Figure 1H, upper right panel**). *Sham* animals maintained the maximal trunk stability score throughout the entire postoperative period. In the *TRI* animal, trunk stability did not fall to zero even at the 1st WPI (**Figure 1H, upper right panel**). Over time, an improvement in trunk stability was observed; its maximal value (“3”) was reached by the 6th WPI (**Figure 1H, upper right panel**).

Altogether, the analysis of sensorimotor functions indicates a direct relationship between the functional disruption and the severity of spinal cord tissue damage revealed by Sudan staining analysis, in which the *TRC2* group had more severe spinal cord damage, especially in its caudal part, than the *TRC1* group.

## 3. Interneuronal connectome recovery

The ability of the injured animals to perform well-coordinated quadrupedal stepping points that the spinal neuronal networks controlling the forelimbs and hindlimbs were coupled in our animals, similar to those in norm (see Frigon, 2017). Previously, Nogueira-Rodrigues et al. (2022) reported fiber regrowth in transected spiny mice. Therefore, we analysed possible connectome recovery in animals of the *TRC1* group by immunostaining for the cytoskeleton protein βIII-tubulin and by the retrograde labeling of possible rewiring of direct connections between the regions below and above the transection using the tracer FB; an advantage of this tracer is the persistence of its labeling, as reported (Novikova et al., 1997).

### 3.1. Damaged area neuronal recovery

In searching for neurogenesis, cells expressing the neuron-specific markers NeuN (Mullen et al., 1992; Sarnat et al., 1998) and βIII-tubulin (Menezes, Luskin, 1994; Kapitein, Hoogenraad, 2015) were assessed within the primary damage region and within the 2-mm secondary damage region (1 mm rostral and 1 mm caudal to the primary damage region). Only cells with DAPI-labeled nuclei were included in the analysis.

#### 3.1.1. βIII-tubulin-labeled processes

In all injured animals, clearly visible βIII-tubulin-labeled fibers were observed both around and within the primary damage region (**Figure 2E, lower left panels**). Typically, βIII-tubulin-labeled fibers were numerous around the primary damage region, especially on its rostral side, but their density decreased drastically within the primary damage region. In the sham-operated and intact animals, only faintly labeled and sparsely distributed βIII-tubulin-labeled fibers and cells were observed. We quantitatively compared βIII-tubulin labeling within the primary damage region across the *TRI*, *TRC1*, and *TRC2* groups (**Figure 2E, lower central diagram**). The *TRI* animal had 9 fibers *per* slice. In the *TRC1* group, there were 4.5-33 fibers *per* slice: the animal that did not exhibit secondary locomotor disruption (#*142*) had a number of labeled fibers comparable to other *TRC1* animals (9.5 *per* slice). In the *TRC2* animals, there were 2.5-21.5 fibers *per* slice; interestingly, the most recovered animal in this group (#*146*) had fewer βIII-tubulin-labeled fibers (2.5 *per* slice). No significant differences were observed between the *TRC1* and *TRC2* groups (U = 6.5, p = 0.2857; M-W).

**Figure 2.**
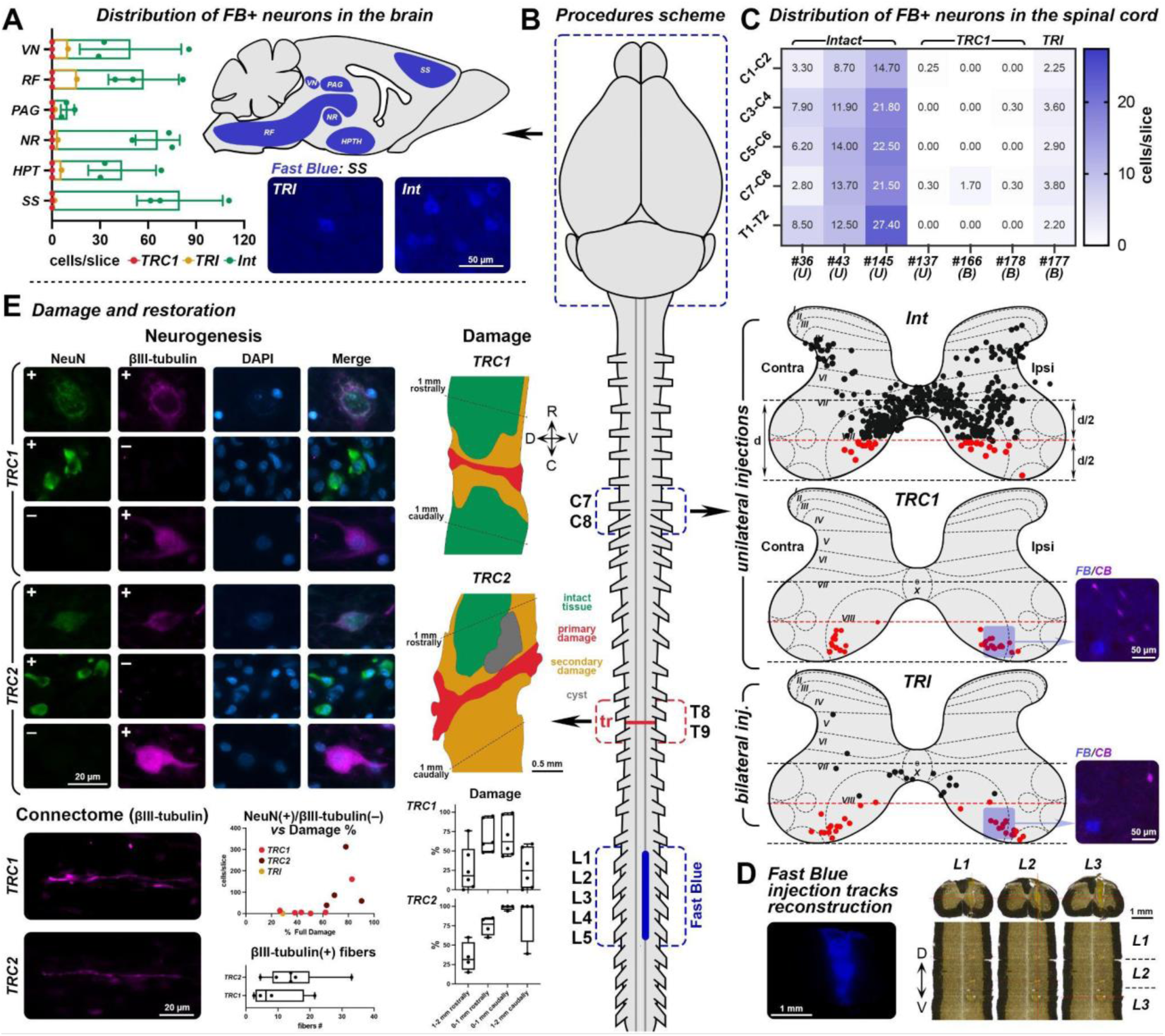
Neuronal recovery in spiny mice after the spinal cord transection. (A) Distribution of Fast Blue-labeled (FB+) neurons in some brain structures: hypothalamus (HPT), nucleus ruber (NR), periaqueductal grey (PAG), reticular formation (RF), somatosensory cortex (SS), and vestibular nuclei (VN), for three groups (regular complete transection, *TRC1*; incomplete transection, *TRI*; intact, *Int*); lower panels – examples of FB+ neurons in the SS cortex of intact (#36) and *TRI* (#177) animals. The scale bar is 50 µm. (B) General scheme of the tracing experiment (transection, tr; cervical, C; thoracic, T; lumbar, L; sacral, S). (C) Distribution of FB+ neurons in the gray matter of the spinal cord: upper panel – heatmap showing the average number of FB+ neurons in rostral spinal segments C1-T2 in *intact*, *TRC1*, and *TRI* animals (bilateral FB injections, B; unilateral FB injections, U); lower panels – schematic view of FB+ neurons (marked by dots) in the gray matter of segments C7-C8 for *intact*, *TRC1*, and *TRI* animals. In the ventral horns, the distance (d) between the central canal and the ventral border of the gray matter was divided into two zones by the red dashed line passing through the midpoint of d; FB+ neurons below the red line are marked in red. To the right of the schemes, enlarged images showing the overlay of FB+ (blue) and calbindin (CB)-labeled (magenta) neurons are displayed. The scale bar is 50 µm. (D) Reconstruction of FB injection tracts (using #36 as an example). Left panel – fluorescent microscope micrograph of the injection track; right panel – light microscope images showing a clear yellow track in transverse slices (upper) and horizontal reconstruction (lower). The scale bar is 1 mm. (E) Neurogenesis in the injury region after spinal cord transection in the *TRC1* and *TRC2* groups. Upper right panel – schematic view of the injured region in two groups. Upper left panel – examples of neuronal somas labeled by the NeuN and βIII-tubulin. Lower left panel – examples of βIII-tubulin-labeled (βIII-tubulin(+)) axons passing through the primary injured region. Lower central diagrams: the relationship of the number of NeuN(+)/βIII-tubulin(–) cells to the total damage percentage in the transection zone (upper), and the number of βIII-tubulin(+) axons (lower); lower right diagrams: the tissue damage percentage along the rostrocaudal axis for the *TRC1* (upper) and *TRC2* (lower) groups.

#### 3.1.2. βIII-tubulin-labeled and NeuN-labeled cells

Based on βIII-tubulin and NeuN expression, three types of neurons were identified: (1) βIII-tubulin(+)/NeuN(+): (2) βIII-tubulin(+)/NeuN(–); (3) βIII-tubulin(–)/NeuN(+) (**Figure 2E, upper left panels**). Anti-βIII-tubulin and anti-NeuN antibodies labelled cell soma with or without proximal processes, respectively. All detected cells were relatively small; in case of βIII-tubulin labeling, cells had a spindle or stellate shape; in case of NeuN labeling, their soma could be described as polygonal.

***βIII-tubulin(+)/NeuN(+)*** cells localized in the secondary damage region and were not observed in the primary damage region or in the fully damaged caudal area of the *TRC2* animals. This was the smallest cell population, with an average of 0-10 cells *per* slice across all analyzed animals and no difference between the *TRC1* and *TRC2* groups (U = 11.5, p = 0.6818; M-W). NeuN(+) labeling of these cells mainly included an immunopositive nucleus and cytoplasm (**Figure 2E, upper left panels**). Given their proximity to the intact tissue, they could be suggested as surviving post-injury neuronal populations.

***βIII-tubulin(+)/NeuN(–)*** cells (**Figure 2E, upper left panels**) were present in both the primary and secondary damage regions, regardless of damage severity or experimental group (*TRC1 vs TRC2*: U = 10.5, p = 0.5606; M-W). Some βIII-tubulin(+)/NeuN(–) cells were found in the primary damage region, making their post-injury generation plausible. Although their number was small, 1-14.5 cells *per* slice, it significantly outnumbered the βIII-tubulin(+)/NeuN(+) cell population (U = 19.5, p = 0.0055; M-W).

***βIII-tubulin(–)/NeuN(+)*** cells mainly had a NeuN-immunopositive cytoplasm and a NeuN-immunonegative nucleus (**Figure 2E, upper left panels**), which was their distinguishing feature; solitary cells had a NeuN(+) nucleus and resembled βIII-tubulin(+)/NeuN(+) cells and the neuronal population of the intact tissue. This cell population showed heterogeneity in number of neurons: (1) the *TRI* animal contained no any βIII-tubulin(–)/NeuN(+) cells, although it had solitary βIII-tubulin(+)/NeuN(+) and βIII-tubulin(+)/NeuN(–) cells; (2) 5/6 of *TRC1* animals contained small numbers of βIII-tubulin(–)/NeuN(+) (1.5-8 cells *per* slice); and (3) 1/6 of TRC1 animals (*#137*; 159 cells *per* slice) together with all 4/4 *TRC2* animals had much higher numbers of βIII-tubulin(–)/NeuN(+) cells (38-314.5 cells *per* slice) (**Figure 2E, lower central diagram**). βIII-tubulin(–)/NeuN(+) cells were found in all damaged regions (caudal, rostral, and primary). In animals with many βIII-tubulin(–)/NeuN(+) cells, they formed local groups with no clear relationship to the primary damage region: only in 2/5 animals, the majority of these cells lied in the primary damage region; in the other 3/5 animals, the majority lied in the caudal damaged area. No difference was observed between the *TRC1* and *TRC2* groups (U = 3.0, p = 0.0667; M-W). Exclusion of the highly deviating *#137* from the *TRC1* group resulted in a significant statistical difference (*TRC1 vs TRC2*: U = 0, p = 0.0159; M-W). Notably, high numbers of βIII-tubulin(-)/NeuN(+) cells were observed only in animals with a total damage area exceeding 60%, which is consistent with the total damage of *#137*, which was 83%.

### 3.2. Propriospinal afferents

To identify propriospinal neurons with axonal projections to the lumbar enlargement, the retrograde tracer FB was injected into the lumbar segments of animals from the *intact*, *TRC1*, and *TRI* groups (**Figure 2A, B, C, D**). In *intact* animals, unilateral FB injection sites occupied segments L1-L3 (#36 and #43) or segments L2-L3 (#145); in all cases, hundreds of labeled neurons were observed within the cervical and thoracic regions above the transection. The average number of labeled cells *per* slice within the cervical enlargement is presented in **Figure 2C** (upper panel). Most of these cells were located within laminae VII and VIII, as has been previously reported for other rodents (Reed et al., 2006; Brockett et al., 2013).

In operated animals, the FB injection site occupied segments L3-L5 (#177: bilateral injections); in the *TRC1* group, segments L2-L5 (#137: unilateral injections) or segments L2-L4 (#166: bilateral injections, #178: bilateral injections). In the *TRC1* and *TRI* animals, a significantly lower number of labeled cells was observed, with the majority located in segments C7-C8 (**Figure 2C**).

We compared the gray matter regions containing these cells: all labeled neurons were overlaid onto the same representative slice averaged for all animals from the same group (**Figure 2С, central panels**). The ventral horns were subdivided into two regions: upper and lower (black and red in **Figure 2С**, respectively), based on a line drawn through the midpoint of the line connecting the central canal and the most ventral border of the gray matter. In *TRC1* animals, 100% of the labeled cells were located within the lower part of the ventral horns (**Figure 2С, central panels**). In contrast, in the *intact* group, only 5.5-6.05% of the labeled cells were located within this area (**Figure 2С, central panels**). In the *TRI* animal with bilaterally spared spinal cord tissue (#177), 64% were located within the lower part (**Figure 2С, central panels**).

Since the labeled region in transected animals corresponds to the area where Renshaw cells are located (Benito-Gonzalez, Alvarez, 2012; Siembab et al., 2016), we compared the pattern of FB-labeled cells with those expressing calbindin, a selective marker for Renshaw cells within this region of the ventral horn in rodents (Carr et al., 1998; Veshchitskii et al., 2025b). However, in all *TRC1* animals, the region containing FB-labelled cells was located below the region containing Renshaw cells; no overlap between FB-labeled and calbindin-labeled cells was observed (**Figure 2С, right panels**).

### 3.3. Supraspinal afferents

In the same animals, brain slices were analysed to identify supraspinal FB-labeled neurons projecting to the lumbar enlargement (**Figure 2A, B**). Brain structures were identified using the mouse brain atlas (https://atlas.brain-map.org/). In *intact* animals, most FB-labeled neurons were observed in the somatosensory cortex, hypothalamic nuclei, red nucleus, periaqueductal gray, reticular nuclei, raphe nuclei, and vestibular nuclei (**Figure 2A**), similar to findings in mice (Liang et al., 2011; Asboth et al., 2018). Notably, in some cases it was difficult to unambiguously assign the labelled cells to either the raphe nuclei or the neighboring reticular formation; consequently, cells located in close proximity to the midline were excluded from the numerical analysis. In *intact* animals, the maximal cell density was observed in the somatosensory cortex, red nucleus, and reticular formation. In contrast, no labeled neurons were found in the brain of the *TRC1* animals. In the *TRI* animal, numerous FB-labeled neurons were observed in the hypothalamic nuclei, reticular formation, and vestibular nuclei, a moderate number of cells was found in the periaqueductal gray, solitary labeled cells were observed in the red nucleus and somatosensory cortex. The density of the labeled neurons in *TRI* animals was much lower than that in the *intact* group (**Figure 2A**).

## 4. Primary culture

The presence of numerous NeuN-expressing cells within the injury region raised the question of whether spiny mice possess any peculiarities regarding neurogenesis. A possible explanation could be related to the differentiation potential of their stem cells. To address this, a primary culture derived from the bone marrow was established. As is well known, under standard conditions, bone marrow mesenchymal stem cells (BM-MSCs) are pluripotent and capable of differentiating into multiple cell lineages (Plikus et al., 2021; Boldyreva et al., 2025). The simplicity of isolation and differentiation of BM-MSCs make them a unique source of different cell types. However, under standard conditions, BM-MSCs do not differentiate into neurons; neuronal differentiation can be induced by specific chemical activators (Yang et al., 2011).

In the present study, we cultivated spiny mouse BM-MSCs without neurogenic activators. Within the zero passage (*P0*), a heterogeneous population of stem cells was observed: myofibroblast-like cells (**Figure 3A, blue arrow**) and blood cells (**Figure 3A, red arrow**) [similar to those described in the literature (Huang et al., 2015; Abdallah et al., 2019)], and an unknown cell type (**Figure 3A, magenta arrow**). This unknown cell type (hereinafter referred to as “ruffles cells”) had a round-shaped soma and clearly visible radial pseudopodia-like processes (**Figure 3B, magenta arrow**). “Ruffle cells” possessed an amoeboid-like morphology, which probably allows them to migrate through myofibroblast-like cells. We were unable to find similar cells in the literature data. On the 9th day, a clear cellular monolayer was observed.

**Figure 3.**
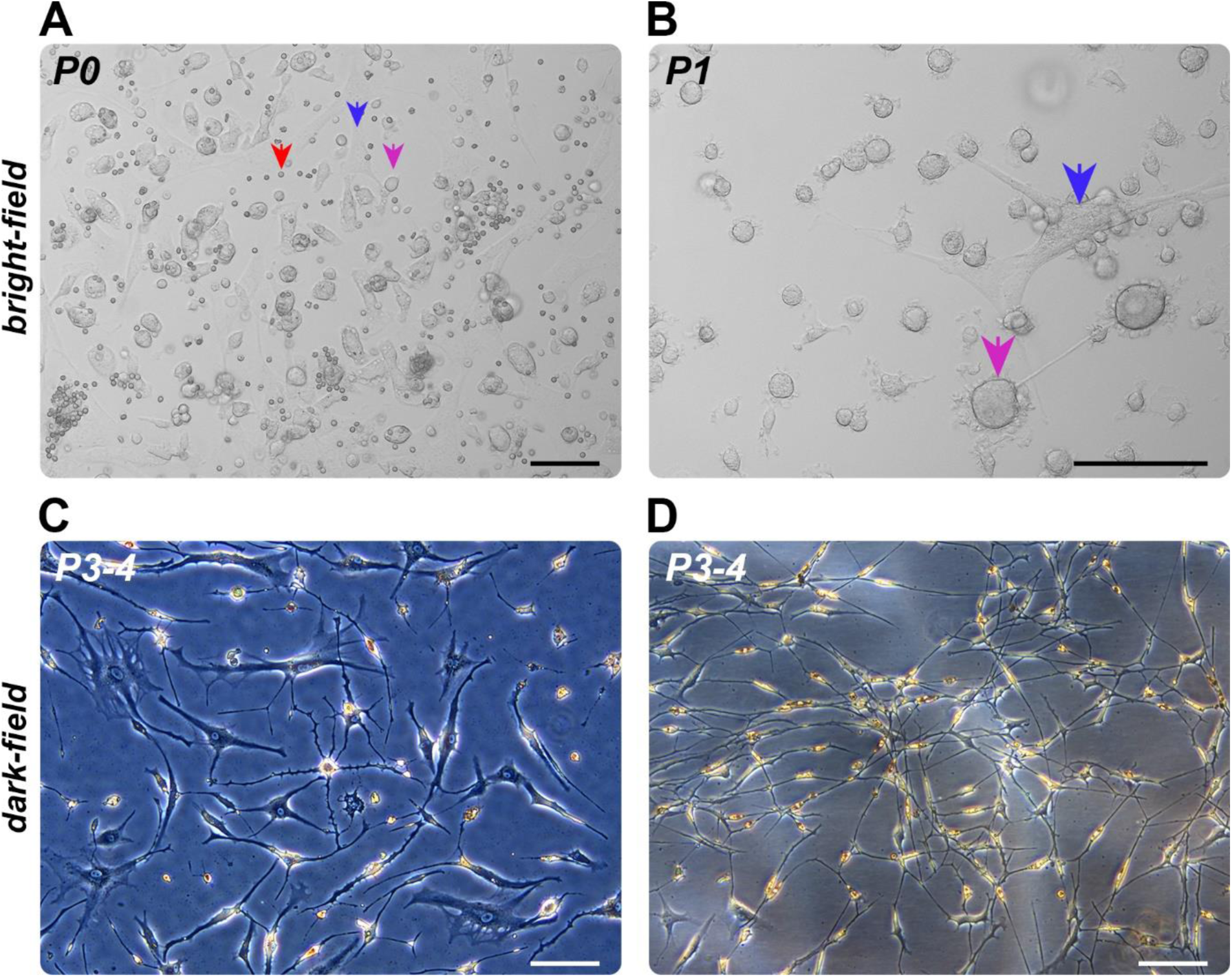
Primary culture of the spiny mice bone marrow mesenchymal stem cells. (A) Passage 0 – myofibroblast-like cells (blue), blood cells (red), and “ruffles cells” (magenta) in bright-field. (B) Passage 1 – myofibroblast-like cells (blue) and “ruffles cells” (magenta) in bright-field. (C) Passages 3-4 – myofibroblast-like and neuron-like cells in dark-field. (D) Passages 3-4 – neuron-like cell population in dark-field.

After the first passage, blood cells completely disappeared; at that stage, the cell colony population consisted of myofibroblast-like cells and “ruffles cells” (**Figure 3B**). The latter were mostly located on the surface of the myofibroblast-like cells. Starting from *P2*, “ruffles cells” completely disappeared; concurrently, transformation of myofibroblast-like cells into neuron-like cells was observed (**Figure 3C**). The primary method to distinguish between fibroblasts and neurons in culture is cell morphology: (1) the neuronal soma is smaller than that of fibroblasts; (2) the cells have thin, branched processes (Vierbuchen et al., 2010; Yang et al., 2011; Samal et al., 2024). Finally, on *P3* and *P4*, on the 7th-10th cultivation days, the culture acquired morphological features similar to those of neuronal cells (**Figure 3D**). Over time, the division rate decreased significantly. Moreover, the neuronal morphology became more pronounced; cellular processes similar to neuronal dendrites covered with spines are illustrated in **Figure 3С**.

In subsequent passages, the cells lost their ability to survive and died. Possible reasons for this include: the medium used may have lacked components critical for maintaining neuronal survival; depletion of growth or trophic factors secreted by precursor cells in earlier passages. An alternative reason for this behavior of the culture may be passivation as a method that exerts mechanical and chemical effects on cells and on the process of their self-differentiation. The exact causes remain to be clarified in further studies.

## 5. Inflammation and astrogliosis

We assessed the integrity of the white matter tracts and gray matter (**Figure 4B**) at different distances from the injury site after spinal cord transection using selective markers of the neuroinflammation and gliogenesis: Iba1 and GFAP. The following spinal regions were analysed: (1) the cervical segment C2, located at the greatest distance from the injury site, (2) the cervical segment C8, the widest segments of the cervical enlargement in the spiny mouse (Veshchitskii et al., 2025b), which contains the labeled propriospinal neurons (see above), (3-4) the thoracic segments T5 and T12, located approximately at equal distances rostral and caudal to the injury site, (5) the lumbar segment L4, which contains the highest number of motoneurons in the lumbar enlargement of the spiny mouse (Veshchitskii et al., 2025a). In the *TRC2* group, it was impossible to section the spinal cord below the transection using a freezing microtome for the histological analysis; therefore, glial detection could not be performed for these animals.

**Figure 4.**
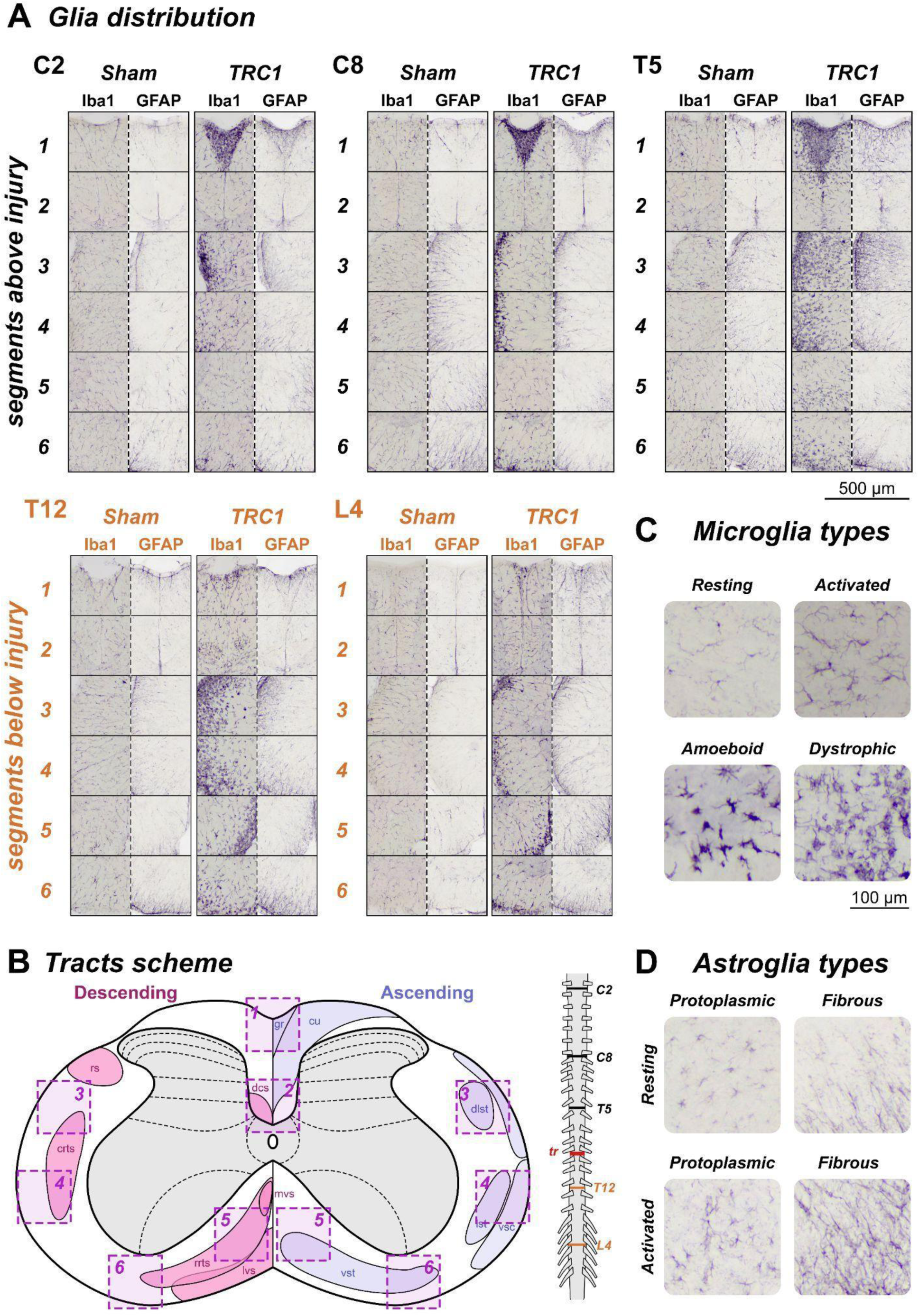
Distribution of microglia (Iba1-labeled cells) and astroglia (GFAP-labeled cells) in spinal cord segments above (C2, C8, and T5) and below (T12 and L4) the injury site in animals from the *TRC1* and *Sham* groups. (A) Distribution of microglia and astroglia in the regions of interest (ROIs) described in B (scale bar: 500 µm.). (B) A scheme of a transverse slice from segment C8 as a representative example with ROIs, and the locations of the analyzed segments on a separate scheme illustrating the rostro-caudal axis of the spinal cord. Presumptive ascending (blue) and descending (pink) tracts in spiny mice (adapted from the Watson, Harrison, 2012) are indicated by the following abbreviations: crts, caudal reticulospinal tract; cu, cuneate fasciculus; dcs, dorsal corticospinal tract; dlst, dorsolateral spinothalamic tract; dsc, dorsal spinocerebellar tract; gr, gracile fasciculus; lst, lateral spinothalamic tract; lvs, lateral vestibulospinal tract; mvs, medial vestibulospinal tract; rrts, rostral reticulospinal tract; rs, rubrospinal tract; vsc, ventral spinocerebellar tract; vst, ventral spinothalamic tract. Higher-magnification images of these ROIs are presented for segments C2, C8, T5, T12, and L4 in A. (C) Examples of microglial phenotypes observed in spiny mice. (D) Examples of astroglial phenotypes observed in spiny mice.

### 5.1. Microglia labeling

Four microglial phenotypes can be distinguished: resting (with small soma size and long ramifications), activated (with increased soma size and shortened ramifications), amoeboid (with rounded soma size and no ramifications), and dystrophic (with fragmented processes) (Lier et al., 2021). Examples of these phenotypes observed in spiny mice during our study are presented in **Figure 4C**.

In the sham-operated animals, mainly solitary activated glial cells were seen in both the gray and white matter. However, solitary clusters of darkly stained glial cells were also observed (**Figure 4A**).

In injured animals, Iba1-labeled cells predominantly exhibited an amoeboid morphology. Within the cervical region, these cells were observed in the gracile fasciculus of the dorsal column and within the lateroventral white matter (**Figure 4A**). Within the thoracic region above the transection, similar staining was additionally seen within the ventromedial white matter (**Figure 4A**). Solitary clusters of darkly stained glial cells were also seen in both the white and gray matter. Below the transection, only weak staining of the gracile fasciculus was observed, along with a sharp increase in staining in the ventral tracts (**Figure 4A**). Within the dorsal corticospinal tract, increased Iba1 labeling was seen below the transection (**Figure 4A**). Notably, no clear differences in Iba1 labeling were observed in animal #*142*, which did not exhibit secondary locomotor impairment (see above).

### 5.2. Astroglia labeling

Resting and activated astroglia were distinguished in multiple spinal cord injury studies. Among resting astroglia, two main types can be observed: protoplasmic astrocytes, located predominantly in the gray matter and characterized by bushy, highly branched processes and irregular cell bodies; and fibrous astrocytes, located predominantly in the white matter and characterized by long, thin, sparsely branched processes (see Sofroniew, 2020). In injured animals, multiple reactive astroglial cells were described, characterized by hypertrophy of the cell body and main processes, thickening of branches, and increased GFAP expression (see Sun, Jakobs, 2012). Examples of these phenotypes observed in spiny mice during our study are presented in **Figure 4D**.

In the sham-operated animals, the typical resting glial distribution was observed in the white matter (Adaes et al., 2017): regardless of the segment, the periphery of the white matter featured a darkly stained network of astroglial processes originating from the GFAP-labeled superficial glia limitans (**Figure 4A**). The network included small, intertwined processes as well as larger processes projecting toward the gray matter, while cell bodies were generally not discernible. A clear zone, an area devoid of dense GFAP-labeled fibers, persisted between the gray matter and the dense network of processes (**Figure 4A**). Individual astrocytes, predominantly elongated toward the gray matter, were observed in this zone.

In injured animals, in segments rostral to the injury site, the gracile tract region was darkly stained (**Figure 4A**). In all segments, the clear zone between the gray matter and the glial process network practically disappeared, with a particularly pronounced glial network developing beneath the dorsal horns, presumably in the region of the rubrospinal tract and dorsolateral spinothalamic tract (**Figure 4A**). Notably, regardless of the segment, the dorsal corticospinal tract region was devoid of a significant number of astroglia, only solitary cells were detected (**Figure 4A**). Note that in animal #*142*, which did not exhibit secondary locomotor impairment, GFAP labeling was similar to that in the sham-operated animals, with the exception of the thoracic segments, where the labeling was similar to that in injured animals.

## DISCUSSION

In a recent paper by Nogueira-Rodrigues et al. (2022), a wide range of stepping abilities in spiny mice after complete spinal cord transection was reported at the 8th WPI: the BMS score for those animals varied from “0” to “8”. This raises the question: are the repair mechanisms after spinal cord transection the same regardless of trauma severity? This issue is extremely important because repair mechanisms after spinal cord injury may depend on injury severity. Based on this, two hypotheses regarding spinal cord regeneration were examined in the present study: (1) recovery of direct descending projections to the lumbar spinal cord, and (2) activation of neurogenesis within the damaged area of the spinal cord.

### Quadrupedal stepping in spinalized spiny mice

In our study, all animals subjected to regular complete transection (*TRC1* group) were able to perform coordinated quadrupedal stepping with weight support. We believe this stepping is coordinated because normal coordination between the fore-and hindlimbs is defined by a one-to-one relationship between their movements (Górska et al., 2013). House mice after complete transection never recover constant coupling between the hindlimbs and forelimbs (Leblond et al., 2003; Nogueira-Rodrigues et al., 2022). In the case of spontaneous recovery of locomotor movements after the complete spinal cord transection in mice, only hindlimb movements, not forelimb movements, were reported (Ung et al., 2007).

Neonate rats and cats after a complete spinal cord transection can spontaneously recover hindlimb locomotion on a treadmill (Bregman, Goldberger, 1983; Commissiong, Sauve, 1993; de Leon et al., 2002). In adult cats, daily training on a treadmill lasting for 2-3 weeks also leads to the gradual recovery of hindlimb locomotion (Rossignol, 2006). In both cases, plastic reorganization of the neuronal networks (due to neonatal plasticity or due to training) is one of the key elements in such restoration. In both cases, cooperation between intrinsic spinal mechanisms and additional sensory stimulation from a moving treadmill was proposed as the main factor driving such recovery (Rossignol, 2006). Notably, in the present study, as previously reported by Nogueira-Rodrigues et al. (2022), no treadmill stimulation was used for spiny mice. It suggests that much weaker sensory stimulation (compared to the treadmill) was efficient for the spiny mice. Furthermore, a high degree of forelimb-hindlimb coordination was observed after spontaneous recovery of stepping following neonatal complete spinal cord transection in opossums; however, a later injury performed at P28 led to an unequal number of forelimb and hindlimb steps (Wheaton et al., 2011, 2013). This finding obviously reflects differences in neuroplasticity between newborn and adolescent animals. Note also that injured animals in our study underwent overground locomotor training three times *per* week. It is possible that adult spiny mice possessed high neuronal plasticity that allows the neuronal connectome re-establishment.

### Interneuronal communication

In the present study, spiny mice from the *TRC1* group developed coordinated quadrupedal stepping around the 4th WPI. It suggests that the functional connections between neuronal networks responsible for the fore-and hindlimbs were restored in injured animals. Nogueira-Rodrigues et al. (2022) reported fiber regrowth in transected spiny mice. We also obtained evidence supporting this feature and analysed the regrowth of intraspinal connections.

Propriospinal connections are among those that facilitate coordination between the forelimbs and hindlimbs (Brockett et al., 2013). Within the spinal cord of transected spiny mice, solitary FB-labeled cells were observed; most of them, if not all, were found in segments C7-C8. Importantly, segments C7-T1 exhibit strong motor rhythmogenic capacity in rats (Ballion et al., 2001) and mice (Gordon et al., 2008). Simultaneous recordings from cervical (C8) and lumbar (L5) ventral roots in rats have revealed coordinated rhythmic activity that alternates between cervical and lumbar levels (Juvin et al., 2012). Interestingly, in opossums subjected to complete spinal cord transection at P7, a control-like pattern of propriospinal cells was observed: at the same time, no labelled cells were found in any spinal segment rostrally to the injury area in animals injured at P28 (Wheaton et al., 2013). These data again highlight the role of neuroplasticity in post-injury repair.

The above-mentioned segments are also interesting because in rodents, lumbar propriospinal neurons terminate mainly within the ventrolateral motor columns located in segments C7-T1 (Matsushita, Ueyama, 1973; Brockett et al., 2013). Given that the location of the fore-and hindlimbs motoneuronal pools within the cervical and lumbar enlargements in spiny mice is similar to those in rats and house mice (Veshchitskii et al., 2025a, 2026), we can conclude that the rhythmogenic neuronal networks in spiny mice have a similar location to those in these rodents. Interestingly, in intact spiny mice, only a few labeled cells were found within the ventral region of segments C7-C8 (most cells were located more dorsally). Therefore, we conclude that propriospinal connections mediated by neurons located in segments C7-C8 were established *de novo*.

Since few direct connections were found in injured spiny mice, we should also keep in mind that in completely transected animals, a repair mechanism involving indirect detour circuits may develop. The cells giving rise to propriospinal interconnections could form these detour circuits, contributing to recovery after partial spinal cord injury in rats (Bareyre et al., 2004; Laliberte et al., 2019). This is supported by the observation that functional recovery after spinal cord transection in mice occurred without maintenance or regeneration of direct supraspinal projections but rather through the reorganization of propriospinal connections (Courtine et al., 2008).

As for the direct supraspinal projections spared in spiny mice after incomplete transection, most FB-labeled cells were found within the hypothalamic nuclei and the reticular formation. Partial sparing of the reticulospinal tracts might be expected, since these tracts are located within the ventrolateral funiculus (Reed et al., 2008). As for the hypothalamic nuclei, in rodents, the hypothalamospinal projection descends through the dorsolateral funiculus (Cechetto, Saper, 1988) and would have been completely destroyed during the spinal transection. It is possible that the hypothalamic tract in spiny mice is located more ventrally, or that hypothalamic projections in animals with incomplete transection were reorganized in this manner. Similarly to the above, we should also keep in mind the possibility of rewiring of supraspinal axons onto propriospinal neurons, consistent with the findings of May et al. (2017), in which rewiring of reticulospinal axons onto lumbar propriospinal neurons was demonstrated.

### Secondary impairment of the quadrupedal stepping

Starting from the 6th WPI, all but one animal from the *TRC1* group (#*142*) exhibited a regression in stepping recovery, manifested as a disruption of the ability to weight support (a switch between the “quadrupedal with support” and “quadrupedal without support” stepping). Possibly, this phenomenon may be related to secondary injury processes, including the formation of cystic cavities. Secondary disruption can result from a plethora of factors, such as vascular damage, excitotoxicity, oxidative stress, inflammation, necrotic cell death, etc (see Alizadeh et al., 2019). Despite the higher resistance of spiny mice (compared to house mice) to oxidative stress and inflammation (Ghebryal et al., 2023), pronounced tissue damage was observed in most animals in the present study.

Secondary injury can lead to muscle spasticity or to a disruption of central balance control. Spasticity is associated with the spontaneous activation of certain motoneuronal pools (Roy, Edgerton, 2012). In rats and rabbits completely spinalized at the thoracic level, muscle spasticity develops gradually (Corleto et al., 2015; Zelenin et al., 2019). The spasticity phenomenon includes an increase in muscle tonus (Marcantoni et al., 2020). However, the secondary locomotor disruption was not accompanied by any change in muscle tonus; therefore, a more complex, non-reflexive mechanism may be proposed.

As for postural control, in experimental mammals, spinalization leads to the disappearance of postural functions (Barbeau et al., 2002; Zelenin et al., 2019) because the ability of spinal networks for postural control, unlike the supraspinal input, is low (Deliagina et al., 2014). After spinal hemisection in neonate cats, the pathways from the brainstem (rubrospinal and vestibulospinal tracts) degenerate (Bregman, Goldberger, 1982). This finding is similar to data obtained in adult cats after complete spinalization, where “…*all vestibulospinal and reticulospinal control of equilibrium has been lost…*” (Rossignol, 2006). At the same time, it is the vestibular system that is implicated in postural adjustments during locomotion (Matsuyama, Drew, 2000). Importantly, throughout recovery, mainly a weak trunk stability was detected in animals with complete transection. Thus, postural control in our experiment remained at moderate-to-low level compared to the high level of leg movements and paw placement.

### Neuroinflammation

Post-injury inflammation includes activation of the resident immune cells (microglia and astrocytes); the maximal activation of these cells occurs during the acute and subacute post-injury phases (see Freyermuth-Trujillo et al., 2022; Bhatt et al., 2024). However, prolonged degenerative processes accompanied by the microglial/macrophagal activation (Wallerian degeneration; see Vargas, Barres, 2007) are evident for more than one month after spinal cord damage (Zhang et al., 1996; Buss et al., 2004). This leads to the so-called “honeycomb-like” appearance of the white matter tracts (Buss et al., 2004). Using Iba1 labeling, we observed that microglial activation persists for at least 70 days after spinal cord injury. Inflammation was detected across multiple segments, mainly in the white matter. Pronounced damage of the dorsal and ventral tracts was observed rostrally to the transection; caudally to the transection, only damage of the ventral tracts was detected. This fact is surprising given the strong downregulation of inflammation in spiny mice (Streeter et al., 2020). At the same time, we observed low GFAP activity; the finding is well consistent with the data of Nogueira-Rodrigues et al. (2022).

Without direct tracing of all ascending and descending spinal tracts, we cannot unambiguously determine the nature of these findings. However, considering a scheme of the major spinal cord tracts in the mouse (Watson, Harrison, 2012), a normal-like Iba1 state can be seen in regions corresponding to the dorsal corticospinal tract and the ventral reticulospinal tract, while increased Iba1 levels were observed in regions corresponding to the dorsal and ventral spinocerebellar tracts, the dorsolateral and lateral spinothalamic tracts, and the cuneatus fasciculus. The gracile fasciculus was damaged only above the transection. The cuneatus fasciculus is entirely located above the transection, in the upper thoracic-cervical region (Watson, Harrison, 2012; Niu et al., 2013), and its integrity was expected. Regarding other ascending and descending spinal tracts, understanding of their integrity is complicated by the fact that the corresponding white matter regions contain multiple tracts. In general, we can speculate that 10 weeks after the spinal cord transection, ascending, rather than descending, spinal tracts were mainly affected. Thus, the observed pattern of damage to ascending spinal tracts with preservation of descending tracts below the injury level represents a classical finding of Wallerian degeneration, in which axons disconnected from their cell bodies undergo progressive degeneration caudal to the lesion site, while segments that retain somatic connections show greater resistance, similar to that observed in other mammals (see Vargas, Barres, 2007).

### Neurogenesis

Multiple cells with markers for neurons (NeuN and βIII-tubulin) were detected within and around the primary injury region. More NeuN-labeled cells were observed in the *TRC2* group than in the *TRC1* group. Notably, the animal in the *TRC1* group with the most damaged tissue had the highest number of these cells, whereas the animal in the *TRC2* group with less damaged tissue had the lowest number. This finding raises the question: is regeneration after the two types of spinal cord transection in the spiny mouse achieved by the same mechanisms or not? It appears that, in contrast to the axonal regrowth observed in both cases, only in the case of particularly severe injury spiny mice are able to enhance neurogenesis.

In this sense, we should note that comparing spiny mice and house mice, over ten times more proliferating cells expressing the cell proliferation marker Ki-67 and twice as many neuroblasts expressing the immature neuron marker doublecortin were detected (Maden et al., 2020). Interestingly, in other precocial rodents, such as degus (*Octodon degus*, *Octodontidae* family) and guinea pig (*Cavia porcellus, Caviidae* family), increased doublecortin expression in the hippocampus has been reported (Akers et al., 2014). Moreover, in degus, widespread doublecortin expression in the cerebral cortex, not restricted to the “classical” neurogenic niches, has been revealed (van Groen et al., 2021). These findings support a higher neurogenic potential in the spiny mouse CNS.

Our data provided evidence for the high proliferation rate of the spiny mice BM-MSCs, together with their fast and almost complete transformation into the neuron-like cells upon the passaging. These findings provide strong evidence for the high ability of spiny mouse stem cells to differentiate into neural cells. In contrast to rat and mouse BM-MSCs, in which only specific chemical stimulation (e.g., lineage-specific transcription factors) can elicit neuronal differentiation (Vierbuchen et al., 2010; Yang et al., 2011; Mu et al., 2015), spiny mouse BM-MSCs were able to differentiate into neurons without any specific stimulation.

Discussing regeneration in spiny mice, a pioneer hypothesis by Seifert et al. (2012) regarding blastema formation (a transient proliferative mass of cells) during ear regeneration should be mentioned, as it resembles the process observed in salamanders and axolotls. Subsequently, several studies have proposed epimorphic regeneration of the ears in spiny mice (Simkin et al., 2017; Seifert, Muneoka, 2018; Avila-Martinez et al., 2023). However, blastema formation in the CNS has not been documented in spiny mice, as in other mammals. At the same time, both salamanders and axolotls are capable of epimorphic regeneration of the spinal cord after the tail amputation (see Fei et al., 2014; Bai et al., 2015; Alibardi, 2019).

Are spiny mice capable of blastema-like formation in the injured CNS? One characteristic of a blastema is histolysis. For example, in house mice, a positive link between the extent of bone lysis and blastema size after the digits amputation has been reported (Seifert, Muneoka, 2018; Dawson et al., 2018), highlighting the “…*relationship between tissue histolysis and progenitor cell availability…*” (Seifert, Muneoka, 2018). Indeed, in our study, as in the study by Kidd et al. (2024), pronounced lysis of the nervous tissue was observed. For example, a progressively increasing percentage of the injured brain region from post-injury day 28 to 168 after middle cerebral artery occlusion was reported (Kidd et al., 2024). Notably, in the *TRC2* group, enormous secondary damage of the spinal cord caudal to the primary injury region was observed. In 3/4 animals, the lumbar region was almost completely lysed, making it impossible to withdraw the tissue for histoprocessing.

We should also discuss the absence of nuclear NeuN labeling within the injured region. NeuN has been identified as a neuron-specific splicing factor, Rbfox3 (Kim et al., 2009), which is a member of the Rbfox family of RNA-binding proteins involving in the regulation of alternative splicing (Kuroyanagi, 2009) and in neuronal development and maturation (Kim et al., 2013; Lin et al., 2016). Although NeuN has been widely used as a common neuronal marker (see Alekseeva et al., 2015), several exceptions have been revealed, including the olfactory bulb mitral cells, Cajal-Retzius cells, cerebellar Purkinje cells, dentate nucleus neurons, etc (Mullen et al., 1992; Sarnat et al., 1998). Moreover, data exist on altered NeuN expression in immature and suffering neurons (Lavezzi et al., 2013). Interestingly, unlike other splicing regulators of this family, Rbfox3 is localized in both the neuronal nucleus and cytoplasm (Lee et al., 2016). Moreover, different variants of the Rbfox3 appear to be predominantly nuclear or cytoplasmic (Dredge, Jensen, 2011). It is possible that NeuN-positive neurons with NeuN-negative nuclei observed here may possess certain features of alternative splicing regulation and/or be damaged, since these cells were mainly detected in animals that showed no repair after severe spinal cord transection.

### Limitations of the study

Regarding the difference in locomotor recovery between male and female in rodents subjected to spinal cord injury, both supporting and contradictory findings exist (Hauben et al., 2002; Swartz et al., 2007; Ung et al., 2007). In any case, in the present study, only females comprised the *TRC1* group, and only males comprised the *TRC2* group.

## RESOURCE AVAILABILITY

### Lead contact

Further information and requests for resources should be directed to the lead contact, Natalia Merkulyeva.

### Materials availability

This study did not generate new unique reagents.

### Data and code availability

Data reported in this paper are available from the lead contact upon request.

## ACKNOWLEDGEMENTS

The study was supported by the funding allocated to the Pavlov Institute of Physiology Russian Academy of Sciences (No: 1021062411653-4-3.1.8). The authors thank the Laboratory of Regulation of Brain Neuronal Functions of the Pavlov Institute of Physiology (chief-lab, E.A. Rybnikova) for the paraffin station; the Laboratory of Experimental Endocrinology of the Pavlov Institute of Physiology (chief-lab, N.I. Yarushkina) and the Institute of Molecular Biology and Genetics of Almazov National Medical Research Center (director, A.A. Kostareva) for providing fluorescent microscopy capacities.

## AUTHOR CONTRIBUTIONS

Conceptualization, NM; data curation, NM; formal analysis, AV, AM, OM, NM; investigation, AV, AM, PS, OG, AB, OM, NM; methodology, AV, AM, PS, OG, AB, OM, AR, NM; project administration, NM; resources, NM, AR; software, AV, AM, OG, NM; supervision, NM; visualization, AV, AM, NM; writing – original draft: AV, AM, PS, OM, NM; writing – review & editing: AV, AM, PS, OG, OM, AR, NM.

## SUPPLEMENTAL INFORMATION

Document S1. Video S1.

## STAR★METHODS

### KEY RESOURCES TABLE

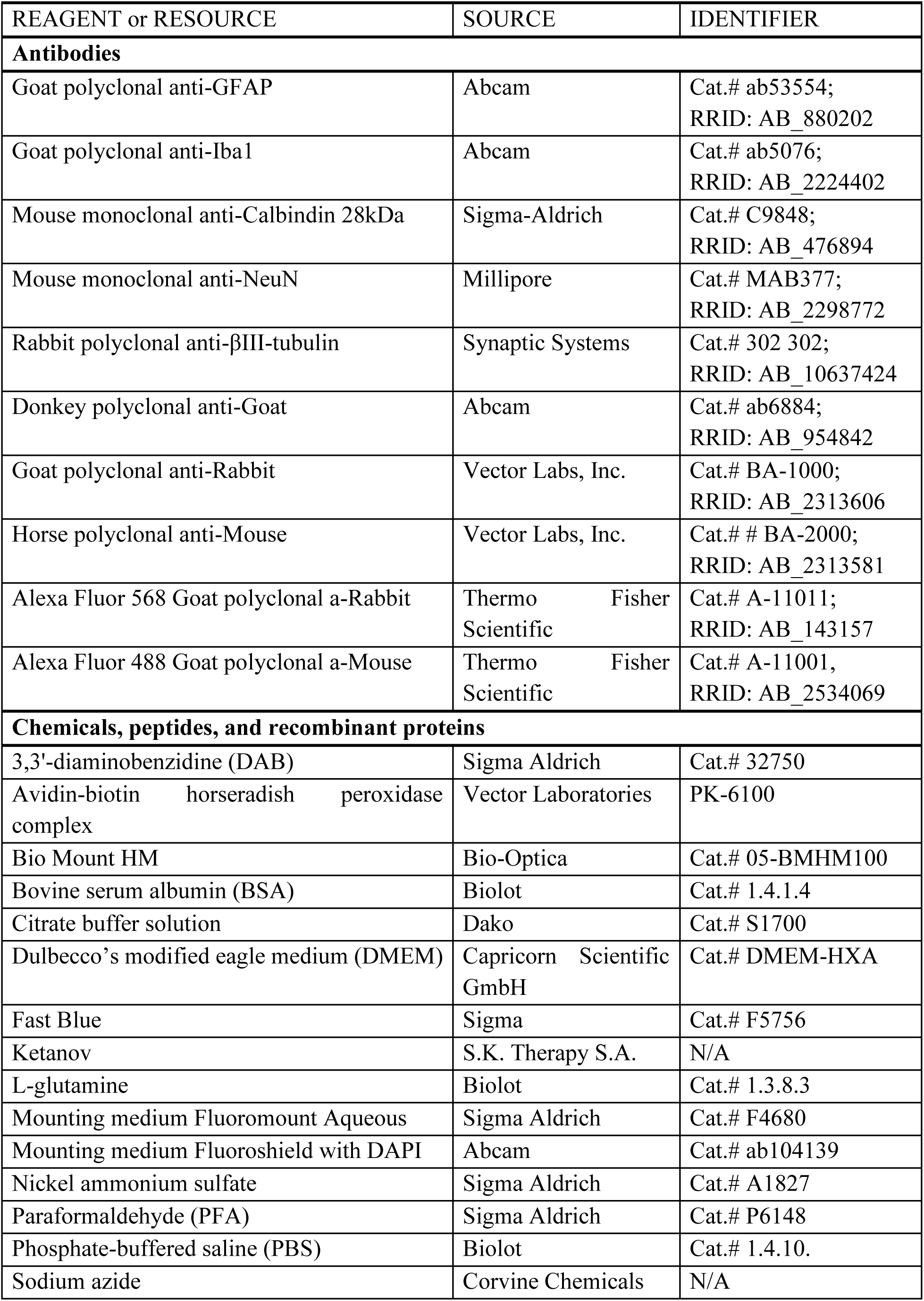
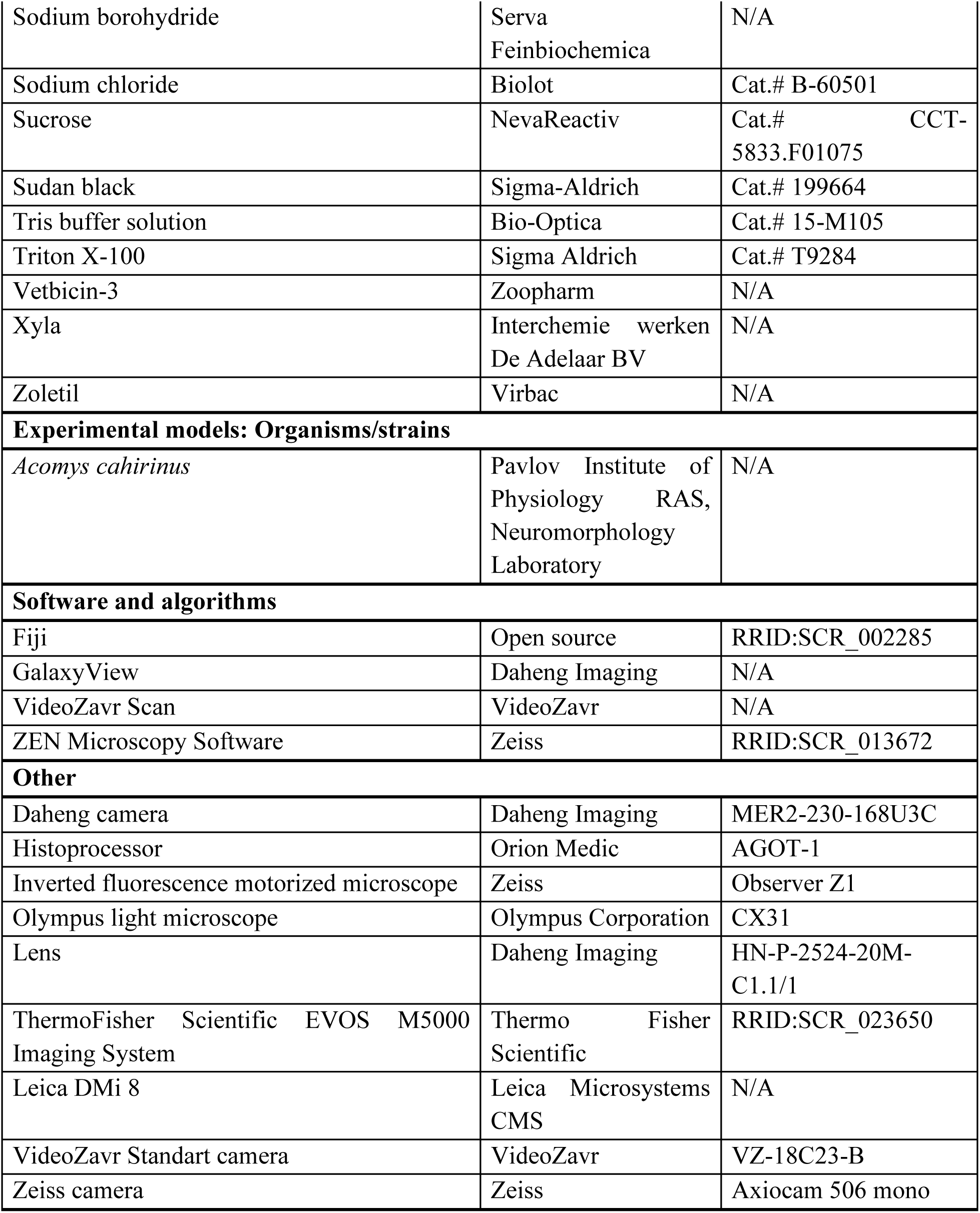

## EXPERIMENTAL MODEL AND STUDY PARTICIPANT DETAILS

### Animals

All experimental procedures were carried out in accordance with the requirements of Council Directive 2010/63/EU of the European Parliament on the protection of animals used for experimental and other scientific purposes and the guidelines of the National Institute of Health Guide for the Care and Use of Laboratory Animals, and with the approval of the Ethics Commission of the Pavlov Institute of Physiology (Protocol #03/14; 14.03.2024). Twenty-five spiny mice (*Acomys cahirinus*) of both sexes, weighing 37-55 g and aged 3-4 months (for the transection experiment) and four spiny mice of both sexes aged 6-12 months (for the primary culture), were used. Animals were bred in the Laboratory of Neuromorphology of the Pavlov Institute of Physiology, housed 3-5 *per* cage, and maintained under standard vivarium light conditions (12 h light/dark cycle) with water provided *ad libitum* and a food diet as elaborated previously (Shkorbatova et al., 2024). Seventeen animals were transected, two animals were used as a sham-operated control, and seven animals were used as an intact control (**Table 1**). The general scheme of the study is presented in **Figure S2**. All animals were handled for 5 days prior to the beginning of the experiments. They were then habituated to the locomotor testing setup and trained to perform the locomotor task (to walk down the track) for 1 week. All animals were also habituated to the urine spot test boxes for 10-14 days before surgery.

## METHOD DETAILS

### Spinal cord transection

All transected animals were subdivided into two groups depending on the method of transection (see below): *TRC1* and *TRC2* (**Table 1**). The completeness of the transection was confirmed *post factum* during the histological analysis. Notably, in the *TRC2* group, enormous secondary damage of the spinal cord caudal to the primary injury site was observed. In most animals, the lumbar region was almost completely lysed, making it impossible to withdraw the tissue for histoprocessing. Accordingly, the *TRC1* group was designated as the “regular transection” group, and the *TRC2* group, as the “severe transection” group.

All surgical procedures were conducted under isoflurane (IsoNik, Chemical Iberica) anesthesia (4% for induction box and 1-2% for maintenance, mixed with air, flow rate 0.8 L/min), administered with Small Animal Anesthesia machine (RWD, R500). During surgery, the animals were placed on a heating pad at 37°C. Under aseptic conditions, a dorsal skin and paravertebral muscle incisions were made in the region above the T7-T12 vertebrae. A complete laminectomy of the T8/T9 vertebra to access the spinal segments T8-T10 according to (Veshchitskii et al., 2025a) and a cut of the *dura mater* were performed, after which the spinal cord was completely transected.

Since the spiny mouse is a new model object in the area of spinal cord injury, two variants for controlling transection completeness were performed: (1) using micro-scissors followed by a microscalpel (Fine Science Tools, 10316-14) – for the *TRC1* group, or (2) using a hook fashioned from half a 26G needle, which was inserted under the spinal cord, after which the spinal cord was gently lifted up; the hook was then used as a guide for the blade of a microscalpel; after transection, the hook was removed through the resulting incision. Both methods have been successfully employed to model spinal cord injury in rats and mice (Leblond et al., 2003; Corleto et al., 2015; Vogelaar, Estrada 2016; Reshamwala et al., 2020); however, the use of micro-scissors alone was described for spine mice (Nogueira-Rodrigues et al., 2022).

Sham-operated animals underwent laminectomy of the T8 vertebra and cut the dura mater without spinal cord damage. The muscles were then sutured with PGA #5 (Lintex, Russia) absorbable sutures, and the skin is sutured with Ethilon #6 (Ethicon, W1615T). After surgery, the animals were placed in a warm recovery cage and monitored until they fully recovered from anesthesia. An antibiotic (Vetbicin-3, Zoopharm; 360 000 U/kg) was administered subcutaneously once after surgery. Analgesia was provided by ketorolac (Ketanov, S.K. Therapy S.A.), with injections (1 mg/kg s/c) twice a day for 3-4 days (Sysoev et al., 2025). Animals were housed in individual cages until the end of the experiment to prevent mutilation of each other.

### Post-injury care

As the transected animals were paraplegic, they were provided with easy access to food and water (hydrogel balls, low-mounted drinkers, and food placed on the floor of the cage) and a high-calorie diet (supplemented with seeds and mealworm larvae). The animals were injected daily with saline (Grotex) until they began to consume adequate amounts of water or hydrogel. Since transection leads to dysfunction of the lower urinary tract, the animals underwent manual bladder emptying in the morning immediately after the urine spot test and before all other tests, and again in the evening until they were able to void voluntarily. Transected animals were carefully monitored daily, with particular attention to the bladder and hindlimbs, until the end of the experiment; the body weight was also checked daily after the testing procedures. If any signs of dehydration, hematuria, and bladder infection appeared, appropriate treatment was administered (saline s/c injection 1-2 ml 2-3 times *per* day, Baytril (Bayer) 2,5% at dose 5-10 mg/kg, diluted in 1 ml of saline, 3-5 days).

Spiny mice that met the criteria for euthanasia (had lost more than 20% of their body weight and showed no trend toward recovery, or exhibited self-mutilation behavior) were euthanized in accordance with humanitarian endpoints guidelines (Lilley et al., 2020; Wang et al., 2025). As a result, six animals were euthanized (three from the *TRC1* group and three from the *TRC2* group) during the first WPI due to self-mutilation with phalangeal autotomy. Moreover, the survival period of animals in the *TRC2* group was two times shorter than that of animals in the *TRC1* group, due to prolonged absence of progression in recovery of locomotor and visceral functions and a persistent body weight loss. Animal care and locomotor testing were performed by four operators. All behavioral scoring and recovery evaluation were conducted by the same two operators.

### Fast Blue injection

FB (Sigma, F5756) was injected into the lumbar spinal cord of four transected animals on the 62-66th post-injury day and in three intact animals. Surgery was carried out under general anaesthesia with isoflurane in air (1.0 L/min flow rate). The initial dose of isoflurane was 5.0%; thereafter, it was maintained at 1.5-2.0%. Under aseptic conditions, a dorsal skin and paravertebral muscle incisions were made in the region above the T11-L1 vertebrae (the peak of spinal kyphosis). A partial laminectomy of the T12 vertebra was performed to access the spinal segments L2-L3 according to (Veshchitskii et al., 2025a). A solution of FB (1.0%, dissolved in distilled water, dH_2O_) was injected unilaterally (intact animals #36, #43, and #145, and transected animal #137) or bilaterally (transected animals #166, #177, and #178) into the spinal cord using a glass pipette (inner diameter: ∼130 µm and outer diameter: ∼450 µm) and a 5-μl syringe (Hamilton Company, PN: 87919) to a depth of 800 μm and 500 μm laterally from the midline at three sites (targeting the central region of the intermediate gray matter according to (Veshchitskii et al., 2025a)), with 0.2 μl *per* injection. After the injection, the glass pipette was left in the tissue for 3 min to avoid tracer leakage. The paravertebral muscles were then sutured with PGA #5 (Lintex) absorbable sutures, and the skin was sutured with Ethilon #6. After surgery, the animal was allowed to recover from anesthesia in a warm box and then replaced in an individual cage. Ketorolac was injected s/c at a dose of 1 mg/kg twice a day for 3 days. The post-injection survival period lasted for 6 days (**Figure S2A**). For spinal cord analysis, every 3rd transverse slice from all spinal segments rostral to the injury site was analyzed. For brain analysis, every 3rd coronal slice was analyzed.

### Evaluation of sensitivity functions

For hindlimb sensitivity tests, the animals were gently secured in a soft cloth, leaving the hindlimbs accessible for testing. Each test was performed starting on the 1st post-injury day, three times *per* week, until the end of the experiment. When the hindlimbs were tested, each was tested individually, but responses were recorded for both the stimulated and the contralateral limb. Since no inter-leg asymmetry was observed, data were averaged across the two hindlimbs. All manual test evaluations were performed by two independent operators who were blinded to each other’s assessments.

#### Muscle tonus evaluation

Each paw was held with two fingers and gently pulled away from and toward the body. Resistance in the limb was evaluated using modified Ashworth Scale, where “0” represented complete limb flaccidity and “5” represented maximal resistance (see details in Yoshizaki et al., 2020).

#### Evaluation of sensitivity to paw pressure

A motor response of the hindlimb was assessed using a manual modification of the Randall-Selitto test (Randall, Selitto, 1957). Each paw was grasped between the thumb and forefinger and gently pressed in the dorsoplantar direction, without forced extension or flexion. The intensity of the withdrawal response was scored on the following 0-5 scale: 0 – no response (absence of any motor reaction even at maximal pressure); 1 – muscle contraction without clear leg movement; 2 – weak withdrawal (partial flexion of the hip); 3 – moderate withdrawal (clear flexion of the hip); 4 – active withdrawal (rapid twitches of the entire limb); 5 – sustained response (prolonged withdrawal, locomotor-like movements).

### Evaluation of sensitivity to pinprick

Motor response to a blunt needle prick was assessed for each hindlimb. Hindlimb motor response to a gentle pinprick on the heel and the ventral surface of each foot was scored on the following 0-5 scale for both the ipsilateral and contralateral limbs: 0 – no response even to strong needle pressure; 1 – slight plantar flexion of the paw in response to strong pressure; 2 – plantar flexion and weak withdrawal of the paw in response to strong pressure; 3 – moderate withdrawal of the paw in response to moderate pressure; 4 – active withdrawal of the leg or 2-3 leg muscle twitches in response to moderate pressure; 5 – prolonged withdrawal of the leg or locomotor-like movements in response to a weak, short touch. The pinprick scale was adapted from Kwiecien et al. (2020) and Ramadhani et al. (2021).

### Urinary function evaluation

Urinary function was tested in two ways: (1) to evaluate bladder contractility, bladder size was assessed *via* abdominal palpation using the following 0-3 scale according to (Streeter et al., 2020): 0 – empty (cannot be palpated); 1 – small; 2 – moderate; 3 – full; (2) to evaluate voluntary micturition function, the urine spot test was performed beginning on the 1st post-injury day (**Figure S2C**). Each animal was placed in an individual box (27 × 17 × 15 cm) with filter paper (NevaReactiv) at the bottom for 30 minutes without access to water and food during the test. Urine spots on the filter paper were visualized under ultraviolet light (**Figure S2C**). This assessment was performed daily for the first 6 weeks and every second day for the last 3 weeks. A calibration curve was generated by applying known volumes of animal urine to filter paper and measuring the resulting spot areas under ultraviolet light. Based on this calibration curve, urine spots were categorized into four size groups: small (<350 mm^2^, corresponding to 30 µl of urine), medium (350-1225 mm^2^), large (> 1225 mm^2^, corresponding to 100 µl of urine), and multiple small drops. The area of each spot was measured using Fiji ImageJ software (open source, ver. 2.16.0/1.54p) (Schindelin et al., 2012), and the number of spots in each category was quantified.

### Behavioral testing

The 0-9 BMS scale (see details in Basso et al., 2006) for locomotion assessment was used. Since the BMS scale does not allow accurate assessment of movements in all joints of the hindlimbs (hip, knee, ankle), which are quite well visible in spiny mice and can provide additional information on functional recovery, especially during the early stages of recovery, we used the corresponding subsection of the BBB test (with a 0-2 scale for each joint; see details in Basso et al., 1995) for this evaluation in a limited number of animals. During a four-minute session, two operators observed the animals’ locomotor behavior in an open field (a box with low walls, 45 × 40 × 3 cm) and completed the BMS and BBB scoring sheets. Since no inter-leg asymmetry was observed, data were averaged across the two hindlimbs.

### Locomotor testing

The experimental walking setup consisted of a glass track (110 × 5 × 15 cm) featuring a non-transparent far wall, and a transparent bottom and near wall (**Figure S2B, left panel**). Light-emitting diodes were positioned alongside the track to ensure uniform illumination of the entire surface. Dark removable “shelters” with a small entrance for the animal were located at each end of the track. A mirror was positioned beneath the entire track at a ∼45-degree angle. A side view and a bottom view (*via* the mirror) were recorded using a camera (Daheng Imaging, MER2-230-168U3C) at 200 frames *per* second. The camera was equipped with a lens (Daheng Imaging, HN-P-2524-20M-C1.1/1) and connected to a computer. Recording was performed using GalaxyView software (Daheng Imaging, ver. 2.6.2107.8081).

Testing began at the 1st WPI and was performed three times *per* week. Animals walked on the track voluntarily or were forced by light touching with fingers or forceps. Videos were analysed manually frame by frame. Only consecutive stepping sequences consisting of at least four steps were analysed. A step cycle (stride) was defined as the time between two sequential steps of the same limb (the left forelimb was chosen as reference one); these moments were detected by footprints on the glass floor (**Figure S2B, right panels**). During analysis, the following six locomotor patterns were identified, and their percentages were calculated: (1) quadrupedal stepping with full or partial body support; (2) quadrupedal stepping without support; (3) tripedal stepping; (4) tripedal stepping with the hindlimb shifted to the next step cycle; (5) bipedal stepping; (6) jumping using the forelimbs for support. All patterns were counted during at least four walking episodes *per* experimental testing. The percentage of the six locomotor patterns was averaged over the 1-week periods. All video analyses were performed by one operator in random order, blinded to the experimental groups.

### Dissection and tissue processing

Animals were deeply anesthetized with a mixture of Zoletil (Virbac; 100 mg/kg; i/p) and Xyla (Interchemie werken “De Adelaar” BV; 20 mg/kg; i/p) and subsequently perfused transcardially with 0.9% sodium chloride (Biolot, B-60501) followed by 4% paraformaldehyde (Sigma Aldrich, P6148). Following perfusion, the spinal cord was removed from the spine, and the brain was removed from the skull, both were then post-fixed overnight. The part of the spinal cord containing the transection area, along with one segment rostrally and one segment caudally, was dehydrated in graded alcohols and mineral oil, infiltrated with paraffin (HistoMix, ErgoProduction) using a histoprocessor (Orion Medic, AGOT-1), and finally embedded in paraffin blocks. Parasagittal slices, 7 μm thick, prepared on a paraffin rotary microtome (Reinhard), were mounted on polysine adhesion slides (SuperFrost) and placed in a thermostat at 37°C for 2-4 weeks to improve adhesion. The remaining spinal cord and brain were placed in 20% and 30% sucrose (NevaReactiv, CCT-5833.F01075) solutions until the tissue sank. The spinal cord was then divided into segments. The rostral edge of each segment was defined as the region containing the caudal-most part of the dorsal rootlet attachment zones (Shkorbatova et al., 2019). Subsequently, each spinal cord segment was cut into 50 μm transverse slices, and the brain was cut into 40 μm coronal slices using a freezing microtome (Reichert).

### Isolation of bone marrow-derived mesenchymal stem cells from spiny mouse

BM-MSCs isolation was carried out under sterile conditions (a second-class laminar flow cabinet with biological protection, in compliance with all biological hazard requirements). The extraction method with modifications was used (Simkin et al., 2024). Animals were rapidly euthanized with CO_2;_ immediately thereafter, the whole body was treated with 70% ethanol by soaking for 2 minutes, after which the femur and tibial bones were collected. To do this, skin, muscles, and tendons were carefully mechanically dissected away from the tibias and femurs using micro-dissecting scissors. The bones were cut at the joints and placed onto a sterile surface (**Figure S2E**). The bone epiphyses were removed, the bone cavity was opened, and the bone marrow was extracted. Notably, this procedure was complicated by the fragility of the spiny mouse bones.

The isolated bones were placed in a sterile culture dish nutrient medium containing Dulbecco’s modified eagle medium (DMEM; Capricorn Scientific GmbH, Ebsdorfergrund-Dreihausen), 10% heat-inactivated fetal bovine serum (FBS), 292 μl/ml L-glutamine (Biolot), and a penicillin-streptomycin mixture (Biolot). Bone marrow was washed using a syringe containing the nutrient medium into a 15-ml centrifuge tube, after which the bone marrow aspirate was centrifuged at 300G for 5 min. The resulting supernatant was discarded, and the cell precipitate was resuspended in 3 ml of DMEM and transferred to T75 flasks for culture. The bone marrow cells were incubated for 9 days in the same medium without medium change; cells were periodically monitored using phase-contrast microscope Leica DMi 8 (Leica Microsystems CMS). On the 9th day, the cells were pressed and their morphology was further observed.

### Passing of bone marrow-derived mesenchymal stem cell culture

On the 9th day, a 100% monolayer of cells was visible; at that moment, cells were washed with DPBS to remove nutrient residues. Then 150 μl of 0.25% trypsin-EDTA (Biolot) was added to the T25 flasks, and cells were incubated for 5 min in an incubator at 37°C in a humid atmosphere containing 5% CO_2._ Trypsin was inactivated by adding the nutrient medium DMEM low glucose (1g/L), 10% FBS, 300 μl L-glutamine, and a penicillin-streptomycin mixture (Biolot). To remove trypsin, detached cells were collected and centrifuged at 500G for 5 min. The cell sediment was decompensated in 1 ml of DMEM and counted using a Garyaev chamber. Thereafter, cells were seeded onto a 24-well plate and into T25 flasks for further culture.

### Injury assessment

The transection degree was assessed based on the analysis of four series (5 slices each) of parasagittal slices taken bilaterally through the lateral and medial levels and spaced equidistantly (**Figure S2D**). Slices were stained with Sudan black (Sigma-Aldrich, cat.# 199664) to reveal lipids, including myelin (Costa et al., 2018). Sudan black staining was also included in the fluorescence immunohistochemistry protocol as a blocker of autofluorescence (see below); images were taken after washing the Sudan black incubation solution from the paraffin slices. Subsequently, the slides were covered with Bio Mount HM (Bio-Optica, cat.# 05-BMHM100) or processed for fluorescence immunohistochemistry. On both unstained and stained slices, the injured tissue was easily determined.

To assess the completeness of the spinal cord transection, a percentage ratio of the damaged area to the intact area within the spinal cord 2 mm caudal and 2 mm rostral to the transection was calculated. The damaged area was subdivided into: (1) the primary damage region – the zone at the transection cut, with fibers oriented along the dissection axis; (2) the secondary damage region – the zone around the primary damage area with signs of gray and white matter destruction. Intact tissue was defined as tissue containing either non-stained neurons of the gray matter or stained neuronal fibers of the white matter, mainly oriented along the spinal cord surface. For each animal, a series of 4 sets of 5 slices each, passing through the mediolateral levels of the spinal cord, were stained with Sudan black, as indicated in **Figure S2D**. This assessment was performed by one operator blinded to the experimental groups.

### Tissue fluorescence immunohistochemistry

To assess the state of the connectome and cytoarchitecture in the injury site and adjacent tissues, fluorescence immunohistochemistry was performed using βIII-tubulin and NeuN antibodies (**Figure S2D**). Slices were processed for immunohistochemistry according to a standard protocol for paraffin-embedded tissue. Paraffin preparations were dewaxed in a series of solvents (xylol ×2, isopropanol 100%, ethanol 96%, 80%, and 70%) for 2-3 minutes each and rehydrated in dH_2O_ ×2 for 5 minutes each. Antigen retrieval was performed in a preheated citrate buffer solution (pH 6.1, Dako, cat.# S1700) with a steamer for 20 minutes. After washing in dH_2O_ ×2 for 5 minutes each, the slices were stained with Sudan black in a concentration of 0.1% in 70% ethanol for 20 minutes at room temperature. This and subsequent incubations were done in a humid chamber. Sudan black staining was adapted from Oliveira et al. (2010) and Yang et al. (2017) and used for both to quench blood autofluorescence at the transection site and as lipophilic staining for histological control of transection quality. Then, slices were washed in phosphate-buffered saline (PBS, 0.01M, pH 7.4, Biolot, cat.# 1.4.10.) ×3 for 5 minutes each. Non-specific antigen binding was blocked with 5% bovine serum albumin (BSA, Biolot, cat.# 1.4.1.4.) for 30 minutes. Thereafter, slices were incubated at 27°C for 72 hours in a solution of PBS with 1% BSA, 0.1% sodium azide (NaN_3;_ Corvine Chemicals) and the following primary antibodies: mouse monoclonal anti-NeuN antibody (1:1000, Millipore, cat.# MAB377, RRID: AB_2298772) or rabbit polyclonal anti-βIII-tubulin antibody (1:500, Synaptic Systems, cat.# 302 302, RRID: AB_10637424). Then, slices were washed in PBS ×3 for 5 minutes each and incubated at 37°C for 2 hours in a solution of PBS with 0.2% Triton X-100 (Sigma Aldrich, cat.# T9284) and the following secondary antibodies: Alexa Fluor 568 goat polyclonal anti-rabbit (1:250, Thermo Fisher Scientific, cat.# A-11011, RRID: AB_143157) or Alexa Fluor 488 goat polyclonal a-mouse (1:250, Thermo Fisher Scientific, cat.# A-11001, RRID: AB_2534069). Then, slices were washed in PBS ×3 for 5 minutes each and coverslipped with DAPI-containing fluorescent mounting medium Fluoroshield (Abcam, cat.# ab104139).

Some slices processed for the FB detection were also labeled for calbindin 28 kDa immunohistochemistry. For this purpose, a standard immunohistochemical protocol on free-floating slices was applied. All following steps were performed under reduced light conditions to preserve the FB labeling. Before the first and between each step, the slices were washed in 0.01M PBS ×3 for 5 minutes each. Antigens retrieval was performed using 1% sodium borohydride (NaBH_4;_ Serva Feinbiochemica). To reduce non-specific binding, slices were incubated in a blocking solution containing 3% BSA. Subsequently, slices were incubated for 70 hours at +4°C in a solution containing the primary mouse monoclonal anti-calbindin 28 kDa antibody (1:3000, Sigma-Aldrich, cat.# C9848, RRID: AB_476894) in PBS with 1% BSA and 0.1% NaN_3._ Following this, slices were incubated for 2 hours at +27°C with Alexa Fluor 488 Goat polyclonal a-Mouse (1:250, Thermo Fisher Scientific, cat.# A-11001, RRID: AB_2534069) in a solution of PBS containing 1% BSA and 0.1% NaN_3._ The slices were then mounted and coverslipped with a mounting medium Fluoromount Aqueous (Sigma, cat.# F4680).

### Tissue DAB immunohistochemistry

To assess the development of microglial activity in the gray and white matter of spinal cord regions located above and below the injury site, Iba1 and GFAP immunohistochemistry was performed on transverse frozen sections using the standard free-floating DAB-immunohistochemical protocol (**Figure S2D**). Before the first and between each step, the slices were washed in 0.01 M PBS ×3 for 5 minutes each. Antigens retrieval was performed using 1% NaBH_4,_ and endogenous peroxidase activity was blocked with 0.3% hydrogen peroxide (H_2O2)_. To reduce non-specific binding, slices were incubated in a blocking solution containing 3% BSA. Subsequently, slices were incubated for 70 hours at +4°C in a solution containing the primary goat polyclonal anti-GFAP antibody (1:1000, Abcam, cat.# ab53554, RRID: AB_880202), or goat polyclonal anti-Iba1 antibody (1:1000, Abcam, cat.# ab5076,RRID: AB_2224402). Following this, slices were incubated for 24 hours at +4°C in a solution containing the donkey polyclonal anti-goat antibody (1:600 Abcam, cat.# ab6884, RRID: AB_954842) in PBS containing 1% BSA and 0.1% NaN_3._ For signal amplification, the avidin-biotin horseradish peroxidase complex (ABC Elite system, Vector Laboratories, Inc., PK-6100) was applied for 1 hour. Staining was visualized using a solution of 1% 3,3’-diaminobenzidine (DAB; Sigma Aldrich, cat.# 32750), 0.1% ammonium nickel(II) sulfate hexahydrate ((NH_4)2N_i(SO_4)2 ·_ 6H_2O_; Sigma Aldrich, cat.# A1827), and 0.03% H_2O2._ Finally, sections were washed in dH_2O_, mounted on slides, air-dried, dehydrated, cleared, and coverslipped with Bio Mount HM.

### Image processing

Images of DAB-and Sudan-stained slices were acquired using an Olympus light microscope (Olympus Corporation, CX31) with a 10× objective, coupled with a VideoZavr Standart camera (VideoZavr, VZ-18C23-B) controlled by VideoZavr Scan software (VideoZavr, ver. 2.4). Images with Alexa-stained slices were acquired using an inverted Zeiss fluorescence motorized microscope (Zeiss, Observer Z1) with 10×, 20×, 63× objectives and a Zeiss camera (Zeiss, Axiocam 506 mono) controlled by Zeiss ZEN (Zeiss, 2.6 blue edition). Images containing FB-and calbindin-labeled neurons were acquired using a fluorescence imaging system (Thermo Fisher Scientific, EVOS M5000). Images of the primary culture were acquired using a phase contrast microscope Leica DMi 8 (Leica Microsystems CMS). Image processing and all measurements were performed using the open-source software Fiji ImageJ.

To identify the spinal segments containing the tracer injection sites, images of unstained wet transverse spinal cord slices from segments L1-L5 were aligned using the central canal and the borders between the white and gray matter as reference points, then stacked and saved as a TIFF file. Using the “Orthogonal Views” function (Ferreira, Rasband, 2012) in Fiji ImageJ, the injection sites were reconstructed in the horizontal and sagittal planes. The injection sites were identified as vertical tracks filled with a yellow FB solution, which was clearly visible in the slices. Subsequently, the presence of the tracer within these tracks was verified under a fluorescence microscope as intensely bright regions.

## QUANTIFICATION AND STATISTICAL ANALYSIS

All parameters measured during each WPI were averaged. Data were analysed using GraphPad Prism software (ver. 8.0.1). Data are presented as mean ± SD. To compare parameters between the *TRC1* and *TRC2* groups, Mann-Whitney U test (M-W) was used.

## SUPPLEMENTAL FIGURES

**Figure S1.**
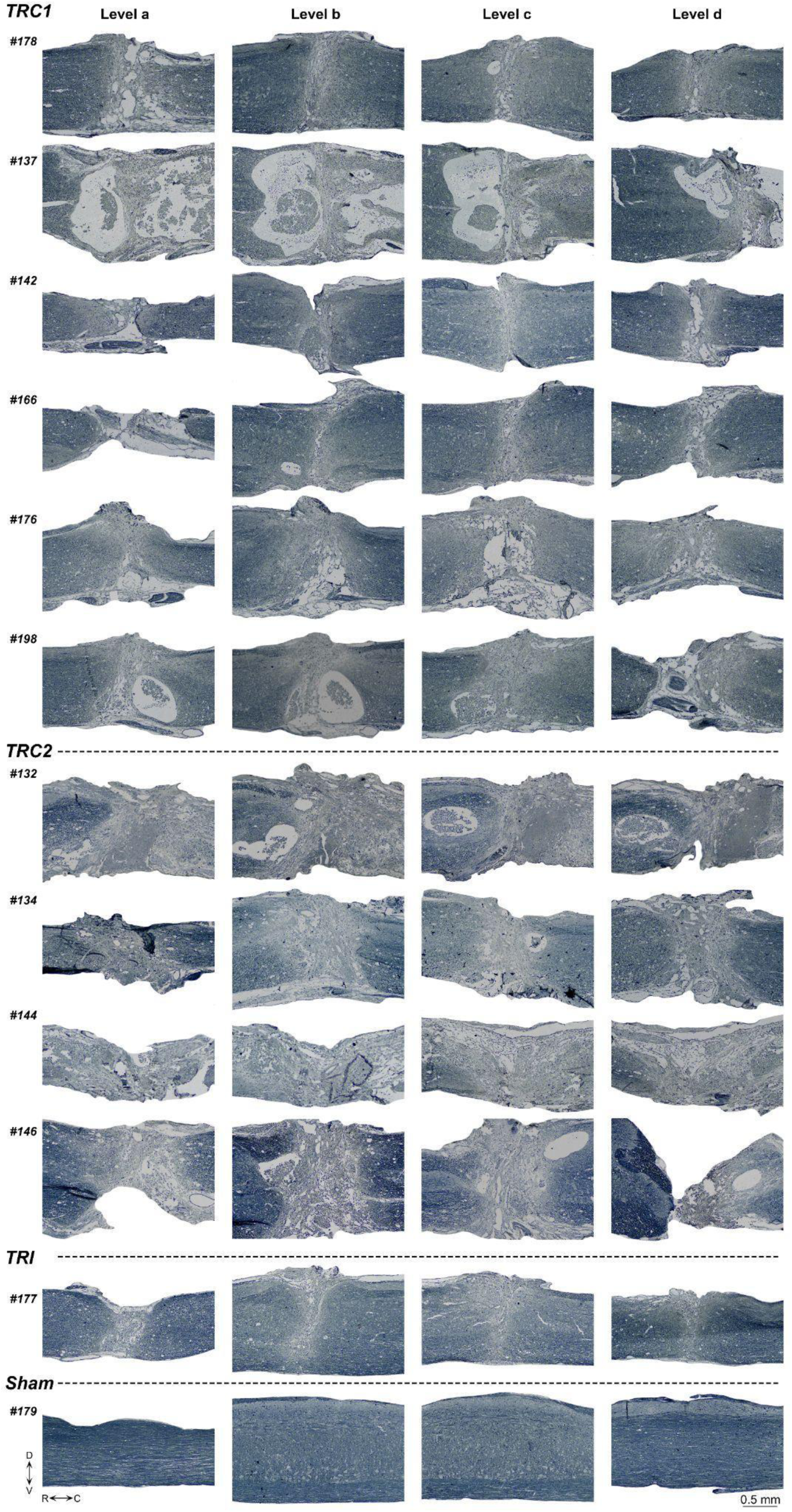
Completeness of the transection analysis. Parasagittal slices stained with Sudan black are shown at four mediolateral levels (a, b, c, and d) of the spinal cord (as indicated on the scheme in **Figure S2D**). *TRC1* – regular complete transection, *TRC2* – severe complete transection. Directions of slice placement are labeled as follows: D – dorsal, V – ventral, R – rostral, C – caudal. WPI – weeks post-injury. The scale bar is 0.5 mm.

**Figure S2.**
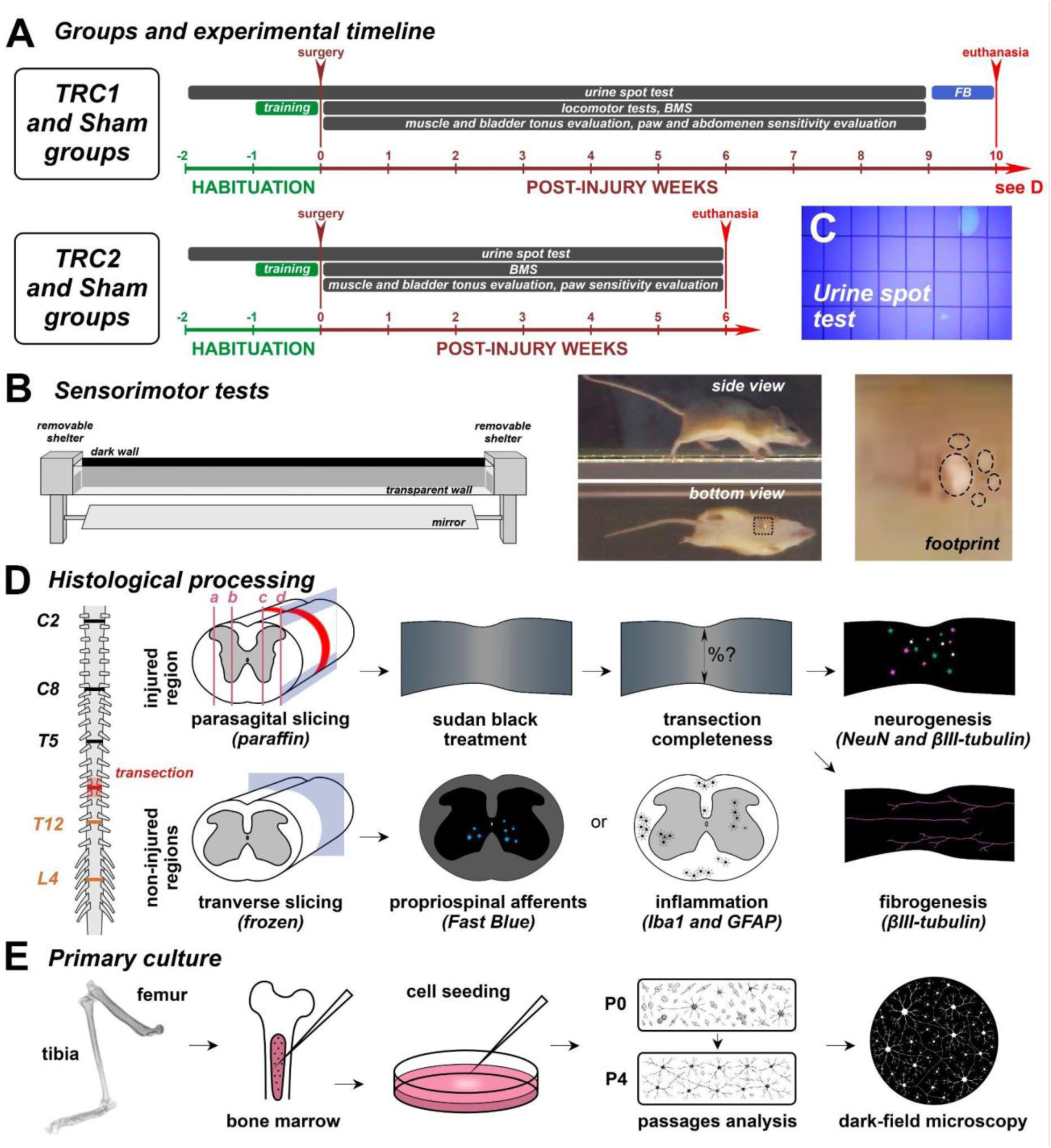
The general scheme of the study. (A) Groups and experimental timeline. Bars indicate the duration of different procedures. (B) Experimental setup for locomotion tests, with an example of side and bottom views from the analyzed recordings, and an example of footprint during walking. (C) An example of filter paper under ultraviolet light after the urine spot test (D) Histological analysis steps for the two types of tissue processing: paraffin for the injured spinal cord part and frozen for the uninjured spinal cord parts. (E) Procedure for the isolation of bone marrow mesenchymal stem cells from the tibia and femur, cell seeding, and primary culture maintenance.

## SUPPLEMENTAL TABLES

**Table S1.** Experimental animals. *TRC1* – regular complete transection, *TRC2* – severe complete transection, *TRI* – incomplete transection, *Sham* – sham-operated. Numericals – individual identifiers of experimental animals.

|  | *40-41 days* | *66-71 days* |
| --- | --- | --- |
| *TRC1* | – | n = 6<br>(#137, #142, #166, #176, #178, #198) |
| *TRC2* | n = 4<br>(#132, #134, #144, #146) | – |
| *TRI* | – | n = 1<br>(#177) |
| *Sham* | n = 1<br>(#135) | n = 1<br>(#179) |
| *Intact* | n = 11<br>(#36, #43, #145, #153, #162, #186, #190, #201, #226, #234, #313) |  |

## SUPPLEMENTARY

## General physiology

### 1.1. Weight loss

Two post-injury stages were compared: the 6th WPI and the 9th WPI. At the 6th WPI, all animals in the *TRC1* group performed moderate weight loss: 3-23%; at the 9th WPI, this weight loss was 5-21%. In the *TRC2* group, moderate to severe weight loss was observed; at the end of the 6th WPI, the weight loss was 19-41%; this was the second reason to terminate the experiment in this group. The first reason was that animals after severe injury chewed their insensate hindlimbs and tail very aggressively. Attempts to stop this behaviour by painting their hindlimbs with various foul-tasting liquids (see Zhang et al., 2001) failed. A similar problem has been described in opossums in (Wheaton et al., 2013). In the *Sham* group, only an increase in body weight was observed: 14-17% at the 6th WPI and 24% at the 9th WPI. In the *TRI* animal, an increase in body weight was observed: 4% at the 6th WPI and 10% at the 9th WPI.

### 1.2. Urinary function

Bladder voiding in the *TRC1* group was detected as early as the 1st WPI, similar to that reported by Nogueira-Rodrigues et al. (2022). In contrast, bladder voiding in the *TRC2* group was worse to that in the *TRC1* group.

All *TRC1* and *TRC2* animals exhibited a high bladder score during the 1st WPI, indicating bladder overfullness and the absence of normal voiding (**Figure 1С**). Over time, the bladder function in *TRC1* animals improved and, starting from the 4th WPI, remained at or below the 0.5 level (**Figure 1С**). In the *TRC2* group, in 3/4 animals, no improvement in bladder function was detected (**Figure 1С**). Only one animal (#*146*) exhibited a decrease in the bladder score beginning at the 3rd WPI. In the *Sham* group, an intact bladder score level was maintained throughout the entire postoperative period (**Figure 1С**). A *TRI* animal performed an intact bladder score starting from the 2nd WPI (**Figure 1С**).

The urine spot test was used to assess urethral sphincter function. In the *TRC1*, *TRI*, and *Sham* animals, no local drops were observed. At the same time, in the *TRC2* group (with the exception of #*146*), numerous small drops were detected throughout the entire postoperative period, with a slight decrease during the last two WPI. As a result, small drops were the predominant type of urine spots for the *TRC2* group. In the *TRC1*, *Sham* and *TRI* animals, predominantly medium-sized spots were observed. The good restoration of urinary function in the *TRC1* group is in good agreement with the findings of Nogueira-Rodrigues et al. (2022).

### 1.3. Sensitivity to pinprick and pressure

Before injury, no hypersensitivity in response to a *hindlimbs paw* pinprick was documented (zero sensitivity). All *TRC1* animals exhibited hypersensitivity to paw prick as early as at the 1st post-injury day. The maximal value of the hypersensitivity (“5.0”) was achieved at the 5th WPI (**Figure 1E**). In the *TRC2* group, only one animal (#*146*) exhibited hypersensitivity to the prick; it achieved the maximal value at the 5th WPI (**Figure 1E**). In the *Sham* group, no hypersensitivity developed (**Figure 1E**). In the *TRI* animal, hypersensitivity to the prick emerged on the 1st postoperative day and never reached the maximal value (**Figure 1E**).

Before injury, no hypersensitivity in response to *paw pressure* was documented (zero sensitivity). In the *TRC1* group, all animals exhibited increased sensitivity to paw pressure as early as the 1st post-injury day (**Figure 1F**). The maximal hypersensitivity score was achieved at the 5th WPI (**Figure 1F**). In the *TRC2* group, only one animal (#*146*) exhibited hypersensitivity to pressure (**Figure 1F**). In the *Sham* group, no hypersensitivity developed (**Figure 1F**). In the *TRI* animal, hypersensitivity to pressure emerged at the 1st WPI and never reached the maximal value (**Figure 1F**).

Hypersensitivity after the spinal cord transection is a result of the allodynia development (Sandkühler, 2008). Hypersensitivity/allodynia is a reason for the central sensitization (Zhang et al., 2005; Grace et al., 2011), and the best model for its study is the complete transection of the spinal cord (Sandkühler, 2008; M’Dahoma et al., 2014). Transection at the thoracic level in rats induced marked hyper-reflexia in the hindpaws and strong mechanical allodynia (Pitcher et al., 2013; M’Dahoma et al., 2014). Notably, in the study by M’Dahoma et al. (2014), mechanical allodynia was detected in the cutaneous territory rostral to the lesion site. In contrast, in the present study, mechanical hyper-reflexia was observed not rostrally but caudally to the lesion site.

In rats, progressive development of allodynia was reported: it began at the 2nd WPI (Yates et al., 2008; Pitcher et al., 2013) and reached its maximal value at the 4th week (Pitcher et al., 2013). In our study, a hyper-reflexia in the hindpaws of spiny mice was first detected at the 1st WPI. The single animal from the *TRC2* performed some rehabilitation and tripedal stepping also exhibited well-developed hypersensitivity, whereas no hypersensitivity was documented in non-rehabilitation animals. This could be explained by the more severe injury of the lumbar spinal cord in these animals and, consequently, disruption of the spinal reflex pathways. Interestingly, in rats subjected to mid-thoracic spinal cord injury of different severity, enhanced mechanical sensitivity was observed only in the case of the most severe injury (Hoschouer et al., 2010). Since hypersensitivity is related to the spinal networks (Lütolf et al., 2022), we suggest that animals with different severities of the spinal cord injury possess different levels of the central sensitization.

## REFERENCES

Abdallah, B.M., Alzahrani, A.M., Abdel-Moneim, A.M., Ditzel, N., and Kassem, M. (2019). A simple and reliable protocol for long-term culture of murine bone marrow stromal (mesenchymal) stem cells that retained their in vitro and in vivo stemness in long-term culture. Biol. Proced. Online 21, 3. 10.1186/s12575-019-0091-3.

Adães, S., Almeida, L., Potes, C.S., Ferreira, A.R., Castro-Lopes, J.M., Ferreira-Gomes, J., and Neto, F.L. (2017). Glial activation in the collagenase model of nociception associated with osteoarthritis. Mol. Pain 13, 1744806916688219. 10.1177/1744806916688219.

Akers, K.G., Martinez-Canabal, A., Restivo, L., Yiu, A.P., De Cristofaro, A., Hsiang, H.-L.L., Wheeler, A.L., Guskjolen, A., Niibori, Y., Shoji, H., et al. (2014). Hippocampal neurogenesis regulates forgetting during adulthood and infancy. Science 344, 598–602. 10.1126/science.1248903.

Alekseeva, O.S., Gusel’nikova, V.V., Beznin, G.V., and Korzhevskii, D.E. (2015). Prospects of the nuclear protein neun application as an index of functional state of the vertebrate nerve cells. Zh. Evol. Biokhim. Fiziol. 51, 313–323.

Alibardi, L. (2019). The regenerating tail blastema of lizards as a model to study organ regeneration and tumor growth regulation in amniotes. Anat. Rec. 302, 1469–1490. 10.1002/ar.24029.

Alizadeh, A., Dyck, S.M., and Karimi-Abdolrezaee, S. (2019). Traumatic spinal cord injury: an overview of pathophysiology, models and acute injury mechanisms. Front. Neurol. 10, 282. 10.3389/fneur.2019.00282.

Asboth, L., Friedli, L., Beauparlant, J., Martinez-Gonzalez, C., Anil, S., Rey, E., Baud, L., Pidpruzhnykova, G., Anderson, M.A., Shkorbatova, P., et al. (2018). Cortico-reticulo-spinal circuit reorganization enables functional recovery after severe spinal cord contusion. Nat. Neurosci. 21, 576–588. 10.1038/s41593-018-0093-5.

Avila-Martinez, N., Gansevoort, M., Verbakel, J., Jayaprakash, H., Araujo, I.M., Vitorino, M., Tiscornia, G., van Kuppevelt, T.H., and Daamen, W.F. (2023). Matrisomal components involved in regenerative wound healing in axolotl and Acomys: implications for biomaterial development. Biomater. Sci. 11, 6060–6081. 10.1039/d3bm00835e.

Bachmann, L.C., Lindau, N.T., Felder, P., and Schwab, M.E. (2014). Sprouting of brainstem-spinal tracts in response to unilateral motor cortex stroke in mice. J. Neurosci. 34, 3378–3389. 10.1523/JNEUROSCI.4384-13.2014.

Bai, X., Wang, Y., Man, L., Zhang, Q., Sun, C., Hu, W., Liu, Y., Liu, M., Gu, X., and Wang, Y. (2015). CD59 mediates cartilage patterning during spontaneous tail regeneration. Sci. Rep. 5, 12798. 10.1038/srep12798.

Ballion, B., Morin, D., and Viala, D. (2001). Forelimb locomotor generators and quadrupedal locomotion in the neonatal rat. Eur. J. Neurosci. 14, 1727–1738. 10.1046/j.0953-816x.2001.01794.x.

Barbeau, H., Fung, J., Leroux, A., and Ladouceur, M. (2002). A review of the adaptability and recovery of locomotion after spinal cord injury. Prog. Brain Res. 137, 9–25. 10.1016/s0079-6123(02)37004-3.

Bareyre, F.M., Kerschensteiner, M., Raineteau, O., Mettenleiter, T.C., Weinmann, O., and Schwab, M.E. (2004). The injured spinal cord spontaneously forms a new intraspinal circuit in adult rats. Nat. Neurosci. 7, 269–277. 10.1038/nn1195.

Basso, D.M., Beattie, M.S., and Bresnahan, J.C. (1995). A sensitive and reliable locomotor rating scale for open field testing in rats. J. Neurotrauma 12, 1–21. 10.1089/neu.1995.12.1.

Basso, D.M., Fisher, L.C., Anderson, A.J., Jakeman, L.B., McTigue, D.M., and Popovich, P.G. (2006). Basso Mouse Scale for locomotion detects differences in recovery after spinal cord injury in five common mouse strains. J. Neurotrauma 23, 635–659. 10.1089/neu.2006.23.635.

Benito-Gonzalez, A., and Alvarez, F.J. (2012). Renshaw cells and Ia inhibitory interneurons are generated at different times from p1 progenitors and differentiate shortly after exiting the cell cycle. J. Neurosci. 32, 1156–1170. 10.1523/JNEUROSCI.3630-12.2012.

Bhatt, M., Sharma, M., and Das, B. (2024). The role of inflammatory cascade and reactive astrogliosis in glial scar formation post-spinal cord injury. Cell. Mol. Neurobiol. 44, 78. 10.1007/s10571-024-01519-9.

Boldyreva, M., Zubkova, E., Trubkina, E., Agareva, M., Michurina, S., Alekseeva, N., Beloglazova, I., Ratner, E., Parfyonova, Y., and Stafeev, I. (2025). Osteogenic shift in the adipose-derived stem cells of *Acomys cahirinus* is linked to impaired adipose tissue self-renewal. Front. Cell Dev. Biol. 13, 1603405. 10.3389/fcell.2025.1603405.

Bregman, B.S., and Goldberger, M.E. (1982). Anatomical plasticity and sparing of function after spinal cord damage in neonatal cats. Science 217, 553–555. 10.1126/science.7089581.

Bregman, B.S., and Goldberger, M.E. (1983). Infant lesion effect: I. Development of motor behavior following neonatal spinal cord damage in cats. Brain Res. 285, 103–117. 10.1016/0165-3806(83)90045-7.

Brockett, E.G., Seenan, P.G., Bannatyne, B.A., and Maxwell, D.J. (2013). Ascending and descending propriospinal pathways between lumbar and cervical segments in the rat: evidence for a substantial ascending excitatory pathway. Neuroscience 240, 83–97. 10.1016/j.neuroscience.2013.02.039.

Buss, A., Brook, G.A., Kakulas, B., Martin, D., Franzen, R., Schoenen, J., Noth, J., and Schmitt, A.B. (2004). Gradual loss of myelin and formation of an astrocytic scar during Wallerian degeneration in the human spinal cord. Brain 127, 34–44. 10.1093/brain/awh001.

Carr, P.A., Alvarez, F.J., Leman, E.A., and Fyffe, R.E. (1998). Calbindin D28k expression in immunohistochemically identified Renshaw cells. Neuroreport 9, 2657–2661. 10.1097/00001756-199808030-00043.

Cechetto, D.F., and Saper, C.B. (1988). Neurochemical organization of the hypothalamic projection to the spinal cord in the rat. J. Comp. Neurol. 272, 579–604. 10.1002/cne.902720410.

Clifford, T., Finkel, Z., Rodriguez, B., Joseph, A., and Cai, L. (2023). Current advancements in spinal cord injury research-glial scar formation and neural regeneration. Cells 12, 853. 10.3390/cells12060853.

Commissiong, J.W., and Sauve, Y. (1993). Neurophysiological basis of functional recovery in the neonatal spinalized rat. Exp. Brain Res. 96, 473–479. 10.1007/BF00234114.

Corleto, J.A., Bravo-Hernández, M., Kamizato, K., Kakinohana, O., Santucci, C., Navarro, M.R., Platoshyn, O., Cizkova, D., Lukacova, N., Taylor, J., et al. (2015). Thoracic 9 spinal transection-induced model of muscle spasticity in the rat: A systematic electrophysiological and histopathological characterization. PLoS One 10, e0144642. 10.1371/journal.pone.0144642.

Costa, P.M. (2018). Staining Protocols. In The handbook of histopathological practices in aquatic environments (Elsevier), pp. 83–117. 10.1016/B978-0-12-812032-3.00004-6.

Courtine, G., Song, B., Roy, R.R., Zhong, H., Herrmann, J.E., Ao, Y., Qi, J., Edgerton, V.R., and Sofroniew, M.V. (2008). Recovery of supraspinal control of stepping via indirect propriospinal relay connections after spinal cord injury. Nat. Med. 14, 69–74. 10.1038/nm1682.

Dawson, L.A., Schanes, P.P., Kim, P., Imholt, F.M., Qureshi, O., Dolan, C.P., Yu, L., Yan, M., Zimmel, K.N., Falck, A.R., et al. (2018). Blastema formation and periosteal ossification in the regenerating adult mouse digit. Wound Repair Regen. 26, 263–273. 10.1111/wrr.12666.

de Leon, R.D., Reinkensmeyer, D.J., Timoszyk, W.K., London, N.J., Roy, R.R., and Edgerton, V.R. (2002). Use of robotics in assessing the adaptive capacity of the rat lumbar spinal cord. Prog. Brain Res. 137, 141–149. 10.1016/s0079-6123(02)37013-4.

Deliagina, T.G., Beloozerova, I.N., Orlovsky, G.N., and Zelenin, P.V. (2014). Contribution of supraspinal systems to generation of automatic postural responses. Front. Integr. Neurosci. 8, 76. 10.3389/fnint.2014.00076.

Dredge, B.K., and Jensen, K.B. (2011). NeuN/Rbfox3 nuclear and cytoplasmic isoforms differentially regulate alternative splicing and nonsense-mediated decay of Rbfox2. PLoS One 6, e21585. 10.1371/journal.pone.0021585.

Fei, J.-F., Schuez, M., Tazaki, A., Taniguchi, Y., Roensch, K., and Tanaka, E.M. (2014). CRISPR-mediated genomic deletion of Sox2 in the axolotl shows a requirement in spinal cord neural stem cell amplification during tail regeneration. Stem Cell Reports 3, 444–459. 10.1016/j.stemcr.2014.06.018.

Ferreira, T. and Rasband, W. (2012) ImageJ user guide. Available online: https://imagej.nih.gov/ij/docs/guide/user-guide.pdf

Freyermuth-Trujillo, X., Segura-Uribe, J.J., Salgado-Ceballos, H., Orozco-Barrios, C.E., and Coyoy-Salgado, A. (2022). Inflammation: a target for treatment in spinal cord injury. Cells 11, 2692. 10.3390/cells11172692.

Frigon, A. (2017). The neural control of interlimb coordination during mammalian locomotion. J. Neurophysiol. 117, 2224–2241. 10.1152/jn.00978.2016.

Ghebryal, L.N., Noshy, M.M., El-Ghor, A.A., and Eissa, S.M. (2023). Comparative analysis of *Acomys cahirinus* and *Mus musculus* responses to genotoxicity, oxidative stress, and inflammation. Sci. Rep. 13, 3989. 10.1038/s41598-023-31143-4.

Gordon, I.T., Dunbar, M.J., Vanneste, K.J., and Whelan, P.J. (2008). Interaction between developing spinal locomotor networks in the neonatal mouse. J. Neurophysiol. 100, 117–128. 10.1152/jn.00829.2007.

Górska, T., Chojnicka-Gittins, B., Majczyński, H., and Zmysłowski, W. (2013). Changes in forelimb-hindlimb coordination after partial spinal lesions of different extent in the rat. Behav. Brain Res. 239, 121–138. 10.1016/j.bbr.2012.10.054.

Grace, P.M., Hutchinson, M.R., Bishop, A., Somogyi, A.A., Mayrhofer, G., and Rolan, P.E. (2011). Adoptive transfer of peripheral immune cells potentiates allodynia in a graded chronic constriction injury model of neuropathic pain. Brain Behav. Immun. 25, 503–513. 10.1016/j.bbi.2010.11.018.

Hauben, E., Mizrahi, T., Agranov, E., and Schwartz, M. (2002). Sexual dimorphism in the spontaneous recovery from spinal cord injury: a gender gap in beneficial autoimmunity? Eur. J. Neurosci. 16, 1731–1740. 10.1046/j.1460-9568.2002.02241.x.

Hoschouer, E.L., Basso, M.D., and Jakeman, L.B. (2010). Aberrant sensory responses are dependent on lesion severity after spinal cord contusion injury in mice. Pain 148, 328–342. 10.1016/j.pain.2009.11.023.

Huang, S., Xu, L., Sun, Y., Wu, T., Wang, K., and Li, G. (2015). An improved protocol for isolation and culture of mesenchymal stem cells from mouse bone marrow. J. Orthop. Translat. 3, 26–33. 10.1016/j.jot.2014.07.005.

Jazayeri, S.B., Maroufi, S.F., Mohammadi, E., Dabbagh Ohadi, M.A., Hagen, E.-M., Chalangari, M., Jazayeri, S.B., Safdarian, M., Zadegan, S.A., Ghodsi, Z., et al. (2023). Incidence of traumatic spinal cord injury worldwide: a systematic review, data integration, and update. World Neurosurg. X 18, 100171. 10.1016/j.wnsx.2023.100171.

Jin, K., Minami, M., Lan, J.Q., Mao, X.O., Batteur, S., Simon, R.P., and Greenberg, D.A. (2001). Neurogenesis in dentate subgranular zone and rostral subventricular zone after focal cerebral ischemia in the rat. Proc. Natl. Acad. Sci. USA 98, 4710–4715. 10.1073/pnas.081011098.

Juvin, L., Le Gal, J.-P., Simmers, J., and Morin, D. (2012). Cervicolumbar coordination in mammalian quadrupedal locomotion: role of spinal thoracic circuitry and limb sensory inputs. J. Neurosci. 32, 953–965. 10.1523/JNEUROSCI.4640-11.2012.

Kapitein, L.C., and Hoogenraad, C.C. (2015). Building the neuronal microtubule cytoskeleton. Neuron 87, 492–506. 10.1016/j.neuron.2015.05.046.

Ke, Y., Chi, L., Xu, R., Luo, C., Gozal, D., and Liu, R. (2006). Early response of endogenous adult neural progenitor cells to acute spinal cord injury in mice. Stem Cells 24, 1011–1019. 10.1634/stemcells.2005-0249.

Kidd, B.M., Varholick, J.A., Tuyn, D.M., Kamat, P.K., Simon, Z.D., Liu, L., Mekler, M.P., Pompilus, M., Bubenik, J.L., Davenport, M.L., et al. (2024). Stroke-induced neuroplasticity in spiny mice in the absence of tissue regeneration. NPJ Regen. Med. 9, 41. 10.1038/s41536-024-00386-8.

Kim, K.K., Adelstein, R.S., and Kawamoto, S. (2009). Identification of neuronal nuclei (NeuN) as Fox-3, a new member of the Fox-1 gene family of splicing factors. J. Biol. Chem. 284, 31052–31061. 10.1074/jbc.M109.052969.

Kim, K.K., Nam, J., Mukouyama, Y.-S., and Kawamoto, S. (2013). Rbfox3-regulated alternative splicing of Numb promotes neuronal differentiation during development. J. Cell Biol. 200, 443–458. 10.1083/jcb.201206146.

Ko, H-Y. (2023) Management and rehabilitation of spinal cord injuries. Springer Singapore. p. 907.

Kuroyanagi, H. (2009). Fox-1 family of RNA-binding proteins. Cell Mol. Life. Sci. 66, 3895– 3907. 10.1007/s00018-009-0120-5.

Kwiecien, J.M., Dabrowski, W., Kwiecien-Delaney, B.J., Kwiecien-Delaney, C.J., Siwicka-Gieroba, D., Yaron, J.R., Zhang, L., Delaney, K.H., and Lucas, A.R. (2020). Neuroprotective effect of subdural infusion of serp-1 in spinal cord trauma. Biomedicines 8, 372. 10.3390/biomedicines8100372.

Laliberte, A.M., Goltash, S., Lalonde, N.R., and Bui, T.V. (2019). Propriospinal neurons: essential elements of locomotor control in the intact and possibly the injured spinal cord. Front. Cell Neurosci. 13, 512. 10.3389/fncel.2019.00512.

Lavezzi, A.M., Corna, M.F., and Matturri, L. (2013). Neuronal nuclear antigen (NeuN): a useful marker of neuronal immaturity in sudden unexplained perinatal death. J. Neurol. Sci. 329, 45–50. 10.1016/j.jns.2013.03.012.

Leblond, H., L’Esperance, M., Orsal, D., and Rossignol, S. (2003). Treadmill locomotion in the intact and spinal mouse. J. Neurosci. 23, 11411–11419. 10.1523/JNEUROSCI.23-36-11411.2003.

Lee, J.-A., Damianov, A., Lin, C.-H., Fontes, M., Parikshak, N.N., Anderson, E.S., Geschwind, D.H., Black, D.L., and Martin, K.C. (2016). Cytoplasmic Rbfox1 regulates the expression of synaptic and autism-related genes. Neuron 89, 113–128. 10.1016/j.neuron.2015.11.025.

Liang, H., Paxinos, G., and Watson, C. (2011). Projections from the brain to the spinal cord in the mouse. Brain Struct. Funct. 215, 159–186. 10.1007/s00429-010-0281-x.

Lier, J., Streit, W.J., and Bechmann, I. (2021). Beyond activation: characterizing microglial functional phenotypes. Cells 10, 2236. 10.3390/cells10092236.

Lilley, E., Andrews, M.R., Bradbury, E.J., Elliott, H., Hawkins, P., Ichiyama, R.M., Keeley, J., Michael-Titus, A.T., Moon, L.D.F., Pluchino, S., et al. (2020). Refining rodent models of spinal cord injury. Exp. Neurol. 328, 113273. 10.1016/j.expneurol.2020.113273.

Lin, Y.-S., Wang, H.-Y., Huang, D.-F., Hsieh, P.-F., Lin, M.-Y., Chou, C.-H., Wu, I.-J., Huang, G.-J., Gau, S.S.-F., and Huang, H.-S. (2016). Neuronal splicing regulator RBFOX3 (NeuN) regulates adult hippocampal neurogenesis and synaptogenesis. PLoS One 11, e0164164. 10.1371/journal.pone.0164164.

Lütolf, R., Rosner, J., Curt, A., and Hubli, M. (2022). Indicators of central sensitization in chronic neuropathic pain after spinal cord injury. Eur. J. Pain 26, 2162–2175. 10.1002/ejp.2028.

M’Dahoma, S., Bourgoin, S., Kayser, V., Barthélémy, S., Chevarin, C., Chali, F., Orsal, D., and Hamon, M. (2014). Spinal cord transection-induced allodynia in rats – behavioral, physiopathological and pharmacological characterization. PLoS One 9, e102027. 10.1371/journal.pone.0102027.

Maas, A.I.R., Menon, D.K., Manley, G.T., Abrams, M., Åkerlund, C., Andelic, N., Aries, M., Bashford, T., Bell, M.J., Bodien, Y.G., et al. (2022). Traumatic brain injury: progress and challenges in prevention, clinical care, and research. Lancet Neurol. 21, 1004–1060. 10.1016/S1474-4422(22)00309-X.

Maden, M., Brant, J.O., Rubiano, A., Sandoval, A.G.W., Simmons, C., Mitchell, R., Collin-Hooper, H., Jacobson, J., Omairi, S., and Patel, K. (2018). Perfect chronic skeletal muscle regeneration in adult spiny mice, *Acomys cahirinus*. Sci. Rep. 8, 8920. 10.1038/s41598-018-27178-7.

Maden, M., Serrano, N., Bermudez, M., and Sandoval, A.G.W. (2021). A profusion of neural stem cells in the brain of the spiny mouse, *Acomys cahirinus*. J. Anat. 238, 1191–1202. 10.1111/joa.13373.

Marcantoni, M., Fuchs, A., Löw, P., Bartsch, D., Kiehn, O., and Bellardita, C. (2020). Early delivery and prolonged treatment with nimodipine prevents the development of spasticity after spinal cord injury in mice. Sci. Transl. Med. 12, eaay0167. 10.1126/scitranslmed.aay0167.

Massey, J.M., Hubscher, C.H., Wagoner, M.R., Decker, J.A., Amps, J., Silver, J., and Onifer, S.M. (2006). Chondroitinase ABC digestion of the perineuronal net promotes functional collateral sprouting in the cuneate nucleus after cervical spinal cord injury. J. Neurosci. 26, 4406–4414. 10.1523/JNEUROSCI.5467-05.2006.

Matsushita, M., and Ueyama, T. (1973). Ventral motor nucleus of the cervical enlargement in some mammals; its specific afferents from the lower cord levels and cytoarchitecture. J. Comp. Neurol. 150, 33–52. 10.1002/cne.901500103.

Matsuyama, K., and Drew, T. (2000). Vestibulospinal and reticulospinal neuronal activity during locomotion in the intact cat. II. Walking on an inclined plane. J. Neurophysiol. 84, 2257–2276. 10.1152/jn.2000.84.5.2257.

May, Z., Fenrich, K.K., Dahlby, J., Batty, N.J., Torres-Espín, A., and Fouad, K. (2017). Following spinal cord injury transected reticulospinal tract axons develop new collateral inputs to spinal interneurons in parallel with locomotor recovery. Neural Plast. 2017, 1932875. 10.1155/2017/1932875.

McDonough, A., Hoang, A.-N., Monterrubio, A.M., Greenhalgh, S., and Martínez-Cerdeño, V. (2013). Compression injury in the mouse spinal cord elicits a specific proliferative response and distinct cell fate acquisition along rostro-caudal and dorso-ventral axes. Neuroscience 254, 1–17. 10.1016/j.neuroscience.2013.09.011.

Menezes, J.R., and Luskin, M.B. (1994). Expression of neuron-specific tubulin defines a novel population in the proliferative layers of the developing telencephalon. J. Neurosci. 14, 5399– 5416. 10.1523/JNEUROSCI.14-09-05399.1994.

Mu, M.W., Zhao, Z.Y., and Li, C.G. (2015). Comparative study of neural differentiation of bone marrow mesenchymal stem cells by different induction methods. Genet. Mol. Res. 14, 14169– 14176. 10.4238/2015.October.29.39.

Mullen, R.J., Buck, C.R., and Smith, A.M. (1992). NeuN, a neuronal specific nuclear protein in vertebrates. Development 116, 201–211. 10.1242/dev.116.1.201.

Niu, J., Ding, L., Li, J.J., Kim, H., Liu, J., Li, H., Moberly, A., Badea, T.C., Duncan, I.D., Son, Y.-J., et al. (2013). Modality-based organization of ascending somatosensory axons in the direct dorsal column pathway. J. Neurosci. 33, 17691–17709. 10.1523/JNEUROSCI.3429-13.2013.

Nogueira-Rodrigues, J., Leite, S.C., Pinto-Costa, R., Sousa, S.C., Luz, L.L., Sintra, M.A., Oliveira, R., Monteiro, A.C., Pinheiro, G.G., Vitorino, M., et al. (2022). Rewired glycosylation activity promotes scarless regeneration and functional recovery in spiny mice after complete spinal cord transection. Dev. Cell 57, 440–450.e7. 10.1016/j.devcel.2021.12.008.

Novikova, L., Novikov, L., and Kellerth, J.O. (1997). Persistent neuronal labeling by retrograde fluorescent tracers: a comparison between Fast Blue, Fluoro-Gold and various dextran conjugates. J. Neurosci. Methods 74, 9–15. 10.1016/s0165-0270(97)02227-9.

Oliveira, V.C., Carrara, R.C.V., Simoes, D.L.C., Saggioro, F.P., Carlotti, C.G., Covas, D.T., and Neder, L. (2010). Sudan Black B treatment reduces autofluorescence and improves resolution of in situ hybridization specific fluorescent signals of brain sections. Histol. Histopathol. 25, 1017– 1024. 10.14670/HH-25.1017.

Peng, H., Shindo, K., Donahue, R.R., Gao, E., Ahern, B.M., Levitan, B.M., Tripathi, H., Powell, D., Noor, A., Elmore, G.A., et al. (2021). Adult spiny mice (Acomys) exhibit endogenous cardiac recovery in response to myocardial infarction. NPJ Regen. Med. 6, 74. 10.1038/s41536-021-00186-4.

Pitcher, G.M., Ritchie, J., and Henry, J.L. (2013). Peripheral neuropathy induces cutaneous hypersensitivity in chronically spinalized rats. Pain Med. 14, 1057–1071. 10.1111/pme.12123.

Plikus, M.V., Wang, X., Sinha, S., Forte, E., Thompson, S.M., Herzog, E.L., Driskell, R.R., Rosenthal, N., Biernaskie, J., and Horsley, V. (2021). Fibroblasts: origins, definitions, and functions in health and disease. Cell 184, 3852–3872. 10.1016/j.cell.2021.06.024.

Plikus, M.V., Wang, X., Sinha, S., Forte, E., Thompson, S.M., Herzog, E.L., Driskell, R.R., Rosenthal, N., Biernaskie, J., and Horsley, V. (2021). Fibroblasts: Origins, definitions, and functions in health and disease. Cell 184, 3852–3872. 10.1016/j.cell.2021.06.024.

Ramadhani, A., Sofro, Z.M., Partadiredja, G.(2021) The effect of oral administration of monosodium glutamate on orofacial pain response and the estimated number of trigeminal ganglion sensory neurons of male Wistar rats. BIO Web of Conferences 41, 05007.

Randall, L.O., and Selitto, J.J. (1957). A method for measurement of analgesic activity on inflamed tissue. Arch. Int. Pharmacodyn. Ther. 111, 409–419.

Reed, W.R., Shum-Siu, A., and Magnuson, D.S.K. (2008). Reticulospinal pathways in the ventrolateral funiculus with terminations in the cervical and lumbar enlargements of the adult rat spinal cord. Neuroscience 151, 505–517. 10.1016/j.neuroscience.2007.10.025.

Reed, W.R., Shum-Siu, A., Onifer, S.M., and Magnuson, D.S.K. (2006). Inter-enlargement pathways in the ventrolateral funiculus of the adult rat spinal cord. Neuroscience 142, 1195– 1207. 10.1016/j.neuroscience.2006.07.017.

Reshamwala, R., Eindorf, T., Shah, M., Smyth, G., Shelper, T., St John, J., and Ekberg, J. (2020). Induction of complete transection-type spinal cord injury in mice. J. Vis. Exp. 10.3791/61131.

Rossignol, S., Barrière, G., Alluin, O., and Frigon, A. (2009). Re-expression of locomotor function after partial spinal cord injury. Physiology 24, 127–139. 10.1152/physiol.00042.2008.

Rossignol, S. (2006). Plasticity of connections underlying locomotor recovery after central and/or peripheral lesions in the adult mammals. Philos. Trans. R. Soc. Lond. B. Biol. Sci. 361, 1647–1671. 10.1098/rstb.2006.1889.

Roy, R.R., and Edgerton, V.R. (2012). Neurobiological perspective of spasticity as occurs after a spinal cord injury. Exp. Neurol. 235, 116–122. 10.1016/j.expneurol.2012.01.017.

Samal, S.K., Sharma, M., and Sarma, J.D. (2024). Isolation and enrichment of major primary neuroglial cells from neonatal mouse brain. Bio Protoc. 14, e4921. 10.21769/BioProtoc.4921.

Sandkühler, J. (2009). Models and mechanisms of hyperalgesia and allodynia. Physiol. Rev. 89, 707–758. 10.1152/physrev.00025.2008.

Sarnat, H.B., Nochlin, D., and Born, D.E. (1998). Neuronal nuclear antigen (NeuN): a marker of neuronal maturation in early human fetal nervous system. Brain Dev. 20, 88–94. 10.1016/s0387-7604(97)00111-3.

Schindelin, J., Arganda-Carreras, I., Frise, E., Kaynig, V., Longair, M., Pietzsch, T., Preibisch, S., Rueden, C., Saalfeld, S., Schmid, B., et al. (2012). Fiji: an open-source platform for biological-image analysis. Nat. Methods 9, 676–682. 10.1038/nmeth.2019.

Seifert, A.W., Kiama, S.G., Seifert, M.G., Goheen, J.R., Palmer, T.M., and Maden, M. (2012). Skin shedding and tissue regeneration in African spiny mice (Acomys). Nature 489, 561–565. 10.1038/nature11499.

Seifert, A.W., and Muneoka, K. (2018). The blastema and epimorphic regeneration in mammals. Dev. Biol. 433, 190–199. 10.1016/j.ydbio.2017.08.007.

Shkorbatova, P.Yu., Veshchitskii, A.A., Mikhalkin, A.A., Nikitina, N.I., Belyaev, A.V., and Merkulyeva, N.S. (2024). Breeding of the Cairo spiny mouse (*Acomys cahirinus*) in laboratory conditions. J Evol Biochem Phys 60, 1347–1362. 10.1134/S0022093024040082.

Shkorbatova, Polina.Y., Lyakhovetskii, V.A., Merkulyeva, N.S., Veshchitskii, A.A., Bazhenova, E.Y., Laurens, J., Pavlova, N.V., and Musienko, P.E. (2019). Prediction algorithm of the cat spinal segments lengths and positions in relation to the vertebrae. Anat. Rec. 302, 1628–1637. 10.1002/ar.24054.

Siembab, V.C., Gomez-Perez, L., Rotterman, T.M., Shneider, N.A., and Alvarez, F.J. (2016). Role of primary afferents in the developmental regulation of motor axon synapse numbers on Renshaw cells. J. Comp. Neurol. 524, 1892–1919. 10.1002/cne.23946.

Simkin, J., Aloysius, A., Adam, M., Safaee, F., Donahue, R.R., Biswas, S., Lakhani, Z., Gensel, J.C., Thybert, D., Potter, S., et al. (2024). Tissue-resident macrophages specifically express Lactotransferrin and Vegfc during ear pinna regeneration in spiny mice. Dev. Cell 59, 496–516.e6. 10.1016/j.devcel.2023.12.017.

Simkin, J., Gawriluk, T.R., Gensel, J.C., and Seifert, A.W. (2017). Macrophages are necessary for epimorphic regeneration in African spiny mice. Elife 6, e24623. 10.7554/eLife.24623.

Sofroniew, M.V. (2020). Astrocyte reactivity: subtypes, states, and functions in cns innate immunity. Trends Immunol. 41, 758–770. 10.1016/j.it.2020.07.004.

Sroga, J.M., Jones, T.B., Kigerl, K.A., McGaughy, V.M., and Popovich, P.G. (2003). Rats and mice exhibit distinct inflammatory reactions after spinal cord injury. J. Comp. Neurol. 462, 223–240. 10.1002/cne.10736.

Stewart, D.C., Serrano, P.N., Rubiano, A., Yokosawa, R., Sandler, J., Mukhtar, M., Brant, J.O., Maden, M., and Simmons, C.S. (2018). Unique behavior of dermal cells from regenerative mammal, the African Spiny Mouse, in response to substrate stiffness. J. Biomech. 81, 149–154. 10.1016/j.jbiomech.2018.10.005.

Streeter, K.A., Sunshine, M.D., Brant, J.O., Sandoval, A.G.W., Maden, M., and Fuller, D.D. (2020). Molecular and histologic outcomes following spinal cord injury in spiny mice, *Acomys cahirinus*. J. Comp. Neurol. 528, 1535–1547. 10.1002/cne.24836.

Sun, D., and Jakobs, T.C. (2012). Structural remodeling of astrocytes in the injured CNS. Neuroscientist 18, 567–588. 10.1177/1073858411423441.

Swartz, K.R., Fee, D.B., Joy, K.M., Roberts, K.N., Sun, S., Scheff, N.N., Wilson, M.E., and Scheff, S.W. (2007). Gender differences in spinal cord injury are not estrogen-dependent. J. Neurotrauma 24, 473–480. 10.1089/neu.2006.0167.

Sysoev, Y.I., Shkorbatova, P.Y., Prikhodko, V.A., Kalinina, D.S., Bazhenova, E.Y., Okovityi, S.V., Bader, M., Alenina, N., Gainetdinov, R.R., and Musienko, P.E. (2025). Central serotonin deficiency impairs recovery of sensorimotor abilities after spinal cord injury in rats. Int. J. Mol. Sci. 26, 2761. 10.3390/ijms26062761.

Telegin, G.B., Chernov, A.S., Malyavina, E.V., Minakov, A.N., Kazakov, V.A., Rodionov, M.V., Belogurov, A.A., and Spallone, A. (2023). CSF – injected contrast medium enhances post-traumatic spinal cord cysts. An experimental study in rats. Eur. Rev. Med. Pharmacol. Sci. 27, 6132–6139. 10.26355/eurrev_202307_32969.

Tran, A.P., Warren, P.M., and Silver, J. (2022). New insights into glial scar formation after spinal cord injury. Cell Tissue Res. 387, 319–336. 10.1007/s00441-021-03477-w.

Ung, R.V., Lapointe, N.P., Tremblay, C., Larouche, A., and Guertin, P.A. (2007). Spontaneous recovery of hindlimb movement in completely spinal cord transected mice: a comparison of assessment methods and conditions. Spinal Cord 45, 367–379. 10.1038/sj.sc.3101970.

van Groen, T., Kadish, I., Popović, N., Caballero Bleda, M., Baño-Otalora, B., Rol, M.A., Madrid, J.A., and Popović, M. (2021). Widespread doublecortin expression in the cerebral cortex of the octodon degus. Front. Neuroanat. 15, 656882. 10.3389/fnana.2021.656882.

Vargas, M.E., and Barres, B.A. (2007). Why is Wallerian degeneration in the CNS so slow? Annu. Rev. Neurosci. 30, 153–179. 10.1146/annurev.neuro.30.051606.094354.

Veshchitskii, A., Shkorbatova, P., and Merkulyeva, N. (2025a). Distribution of the motoneuronal pools controlling the hindlimb muscles in the lumbar spinal cord of the *Acomys cahirinus*. Brain Struct. Funct. 230, 86. 10.1007/s00429-025-02946-0.

Veshchitskii, A., Shkorbatova, P., and Merkulyeva, N. (2025b). Neuroanatomical and neurochemical atlas of the spiny mouse (*Acomys cahirinus*) spinal cord. Brain Struct. Funct. 230, 124. 10.1007/s00429-025-02982-w.

Veshchitskii, A., Shkorbatova, P., and Merkulyeva, N. (2026). Distribution of the motoneuronal pools controlling the forelimb muscles in the cervical spinal cord of the *Acomys cahirinus*. Anat. Rec. 309, 1368–1385. 10.1002/ar.70096.

Vessal, M., Aycock, A., Garton, M.T., Ciferri, M., and Darian-Smith, C. (2007). Adult neurogenesis in primate and rodent spinal cord: comparing a cervical dorsal rhizotomy with a dorsal column transection. Eur. J. Neurosci. 26, 2777–2794. 10.1111/j.1460-9568.2007.05871.x.

Vierbuchen, T., Ostermeier, A., Pang, Z.P., Kokubu, Y., Südhof, T.C., and Wernig, M. (2010). Direct conversion of fibroblasts to functional neurons by defined factors. Nature 463, 1035– 1041. 10.1038/nature08797.

Vogelaar, C.F., and Estrada, V. (2016). Eexperimental spinal cord injury models in rodents: anatomical correlations and assessment of motor recovery. In Recovery of Motor Function Following Spinal Cord Injury, H. Fuller and M. Gates, eds. (InTech). 10.5772/62947.

Wang, S., Cai, S., Zhu, Z., Zeng, W., Hu, S., and Shi, B. (2025). Spinal cord injury modeling: from modeling to evaluation using rats as examples. Front. Neurol. 16, 1573779. 10.3389/fneur.2025.1573779.

Wang, Y., Wu, W., Wu, X., Sun, Y., Zhang, Y.P., Deng, L.-X., Walker, M.J., Qu, W., Chen, C., Liu, N.-K., et al. (2018). Remodeling of lumbar motor circuitry remote to a thoracic spinal cord injury promotes locomotor recovery. Elife 7, e39016. 10.7554/eLife.39016.

Watson, C., and Harrison, M. (2012). The location of the major ascending and descending spinal cord tracts in all spinal cord segments in the mouse: actual and extrapolated. Anat. Rec. 295, 1692–1697. 10.1002/ar.22549.

Wheaton, B.J., Callaway, J.K., Ek, C.J., Dziegielewska, K.M., and Saunders, N.R. (2011). Spontaneous development of full weight-supported stepping after complete spinal cord transection in the neonatal opossum, *Monodelphis domestica*. PLoS One 6, e26826. 10.1371/journal.pone.0026826.

Wheaton, B.J., Noor, N.M., Whish, S.C., Truettner, J.S., Dietrich, W.D., Zhang, M., Crack, P.J., Dziegielewska, K.M., and Saunders, N.R. (2013). Weight-bearing locomotion in the developing opossum, Monodelphis domestica following spinal transection: remodeling of neuronal circuits caudal to lesion. PLoS One 8, e71181. 10.1371/journal.pone.0071181.

Yang, J., Yang, F., Campos, L.S., Mansfield, W., Skelton, H., Hooks, Y., and Liu, P. (2017). Quenching autofluorescence in tissue immunofluorescence. Wellcome Open Res. 2, 79. 10.12688/wellcomeopenres.12251.1.

Yang, N., Ng, Y.H., Pang, Z.P., Südhof, T.C., and Wernig, M. (2011). Induced neuronal cells: how to make and define a neuron. Cell Stem Cell 9, 517–525. 10.1016/j.stem.2011.11.015.

Yates, C., Charlesworth, A., Allen, S.R., Reese, N.B., Skinner, R.D., and Garcia-Rill, E. (2008). The onset of hyperreflexia in the rat following complete spinal cord transection. Spinal Cord 46, 798–803. 10.1038/sc.2008.49.

Yoon, J.H., Cho, K., Garrett, T.J., Finch, P., and Maden, M. (2020). Comparative proteomic analysis in scar-free skin regeneration in *Acomys cahirinus* and scarring *Mus musculus*. Sci. Rep. 10, 166. 10.1038/s41598-019-56823-y.

Yoshizaki, S., Yokota, K., Kubota, K., Saito, T., Tanaka, M., Konno, D.-J., Maeda, T., Matsumoto, Y., Nakashima, Y., and Okada, S. (2020). The beneficial aspects of spasticity in relation to ambulatory ability in mice with spinal cord injury. Spinal Cord 58, 537–543. 10.1038/s41393-019-0395-9.

Zelenin, P.V., Lyalka, V.F., Orlovsky, G.N., and Deliagina, T.G. (2019). Changes in activity of spinal postural networks at different time points after spinalization. Front. Cell Neurosci. 13, 387. 10.3389/fncel.2019.00387.

Zhang, H., Xie, W., and Xie, Y. (2005). Spinal cord injury triggers sensitization of wide dynamic range dorsal horn neurons in segments rostral to the injury. Brain Res. 1055, 103–110. 10.1016/j.brainres.2005.06.072.

Zhang, Y.P., Onifer, S.M., Burke, D.A., and Shields, C.B. (2001). A topical mixture for preventing, abolishing, and treating autophagia and self-mutilation in laboratory rats. Contemp. Top. Lab. Anim. Sci. 40, 35–36.

Zhang, Z., Fujiki, M., Guth, L., and Steward, O. (1996). Genetic influences on cellular reactions to spinal cord injury: a wound-healing response present in normal mice is impaired in mice carrying a mutation (WldS) that causes delayed Wallerian degeneration. J. Comp. Neurol. 371, 485–495. 10.1002/(SICI)1096-9861(19960729)371:3%253C485::AID-CNE10%253E3.0.CO;2-I.

